# A mechanistic model in sheep incorporating energy allocation trade-offs between the ewe and her lambs

**DOI:** 10.1101/2025.04.23.650166

**Authors:** M. Hiltpold, R. Trépos, F. Douhard

**Author notes:** Corresponding author: Frédéric Douhard.

## Abstract

Being able to predict the animal’s multiple traits responses to its environment is of high interest to study livestock adaptation to future conditions. In particular, trade-offs between traits are assumed to be a key determinant of animal robustness. Mechanistic models of energy allocation have been developed to predict potential trade-offs due to the competitive use of energy between biological functions, both within an individual’s life and among individuals. However, previous models have not explicitly integrated potential trade-offs between the mother and her offspring as well as among siblings. Yet those features are essential, particularly in suckling systems. Here we present an energy allocation model in sheep that incorporates the energy transfers between the mother and her offspring. The model simulates the trajectories of responses to feeding and herd management during the sheep’s lifetime. We first provide a detailed description of the model following a standard protocol. We then show that simulation results are in a realistic range in general. We finally illustrate how the model can represent individual variation in ewe-lamb trade-offs by varying the value of five input parameters controlling energy acquisition and allocation. Such parameterization allows to define different individual “types” of energy allocation strategies that are characterized by repeatable patterns of ewe and lamb output traits. This new sheep model provides the basis for future individual-based studies.

**Implications:** Addressing animal robustness in future environments is challenging due to the limited data availability on how multiple traits respond to constraining conditions. In this context, the sheep model presented here serves as a valuable tool for predicting the responses of various traits and their potential trade-offs throughout an animal’s lifetime. Its key innovation lies in incorporating trade-offs between the ewe and her lambs, as well as among lambs within a litter. This feature makes the model a strong basis for characterizing robust sheep in suckling systems.

## Introduction

Mechanistic animal models can represent valuable tools for predicting how animals will respond to future environmental conditions. By condensing generalized knowledge from experimental findings or first principles and translating it into a system of mathematical equations, mechanistic models usually have a broad range of applications compared with statistical models. This includes the possibility to predict responses to new farming conditions, for instance as a result of climate change (Black, 2014). Another important application is the possibility to predict the responses of various individuals that make up a flock or a population. For instance, the use of a mechanistic animal model in individual-based simulations enables to explore responses to herd management or genetic selection (e.g. Puillet et al., 2021; Douhard et al., 2014).

Mechanistic models have been applied to specific aspects of animal production such as feed intake (Pulina et al., 2013; Hackmann and Spain, 2010), growth (Tedeschi et al., 2004) and lactation performance in cattle (Beever et al., 1991). Other models have sought to predict the animal’s manifold responses to nutrition, and more specifically the potential trade-offs that might occur between those responses. This aspect is of increasing interest as the animal’s ability to manage trade-offs when facing nutrient scarcity is assumed to be a strong determinant of animal robustness (Friggens et al., 2017).

Trade-offs assumptions are often based on the principle of nutrient partitioning (Friggens and Newbold, 2007), that is commonly referred to as “resource allocation” in ecological models (Sibly et al., 2013). In most cases, trade-offs are addressed within the scope of energy budgets only. Dynamic energy allocation models predicting the entire life trajectory are especially relevant to study trade-offs consequences between traits expressed at different timescales within an individual’s life. In ruminant livestock, many models have focused on dairy animals and trade-offs that may occur between milk production, body reserves, fertility and longevity. Such models were designed for dairy goats (Douhard et al., 2014) and dairy cows (Puillet et al., 2016) and thus primarily focused on lactating females that do not raise their offspring, omitting the nutritional link between mother and offspring. In contrast, trade-offs involving both maternal and offspring traits have received less attention, whereas they likely are critical in suckling systems, notably in polytocous species such as sheep where mothers can give birth and raise multiple offspring at once. A ruminant model that explicitly integrates the energy transfers from the mother to its offspring remains to be developed to address potential mother-offspring trade-offs.

Here we present a mechanistic model of energy allocation in suckler sheep. The model explicitly incorporates the energy transfers from the ewe to its offspring (one or multiple lambs) from conception to weaning, allowing to study ewe-lamb trade-offs. Our aim here is to highlight two key features of our model that will enable the implementation of individual-based simulations. Specifically, ewe and lambs are modelled as separate individuals who are interacting. Moreover, by varying model input parameters we show how our model can simulate individual variability. In the following, we first provide a thorough description of the model following the standard Overview Design Details (ODD) protocol (Grimm et al., 2006, 2020). The ODD protocol makes it easier to read and write individual-based models, and facilitate their replication (Grimm et al., 2020). It includes 7 main sections that are divided in Overview (Purpose and patterns, Entities, State variables and scales, Process overview and scheduling), Design (Design concepts), and Details (Initialization, Input data, Sub-models).

## Material and methods

### Model description

#### 1. Model purpose and patterns

The purpose of our model is to predict the trajectories of trait responses of a suckler ewe and her lambs to feed energy availability and flock management practices. The ultimate purpose of the model is to explore how energy-allocation trade-offs may translate into trade-offs among individuals, especially in populations under selection. Taking this perspective, we first show in this paper how individual variation in energy acquisition and allocation strategies translate into repeatable patterns in ewe and lamb traits. The model should be able to realistically describe the responses of ewe and her lambs as lifetime trajectories of multiple traits. Specifically, traits include feed intake, body masses, body fat reserve proportion, litter size, milk production, lamb growth rate, reproductive and mortality rates, and lifespan.

#### 2. Model entities, state variables and scales

The model entity is the individual sheep. Its attributes are either static over the sheep’s life (e.g., *ID, sex, MOTH*) or dynamic and thus changing over time according to individual’s responses to its environment (e.g., *PropRes, MassTotal*) (Table 1). Dynamic attributes include three main types of elements characterizing the trajectories of responses of an individual: (1) the physiological state, including status (e.g. *Alivestat, Cycling*_*stat*_, *Lact*_*stat*_) and times (e.g. *age, Cycling*_*time*_, *Lact*_*time*_), (2) different flows (e.g. *Acq*_*solid*_) and stocks (e.g. *MassTotal, PropRes*) of matter or energy describing the transformation of feed intake into animal production, and (3) for mothers only, the current number of dependent offspring (e.g. *NLB, NLS*). The model’s timescale is in days. The simulation time step is discrete (one day). The individual’s dynamic attributes are updated every day and saved as model output at each recording time step (*dT*) until the end of simulation (*TSIM*) is reached. Both *dT* and *TSIM* are defined by the user. We suggest to set *dT* at 7 days as this reduces the calculation time while maintaining the possibility to precisely describe trajectories.

**Table 1:**
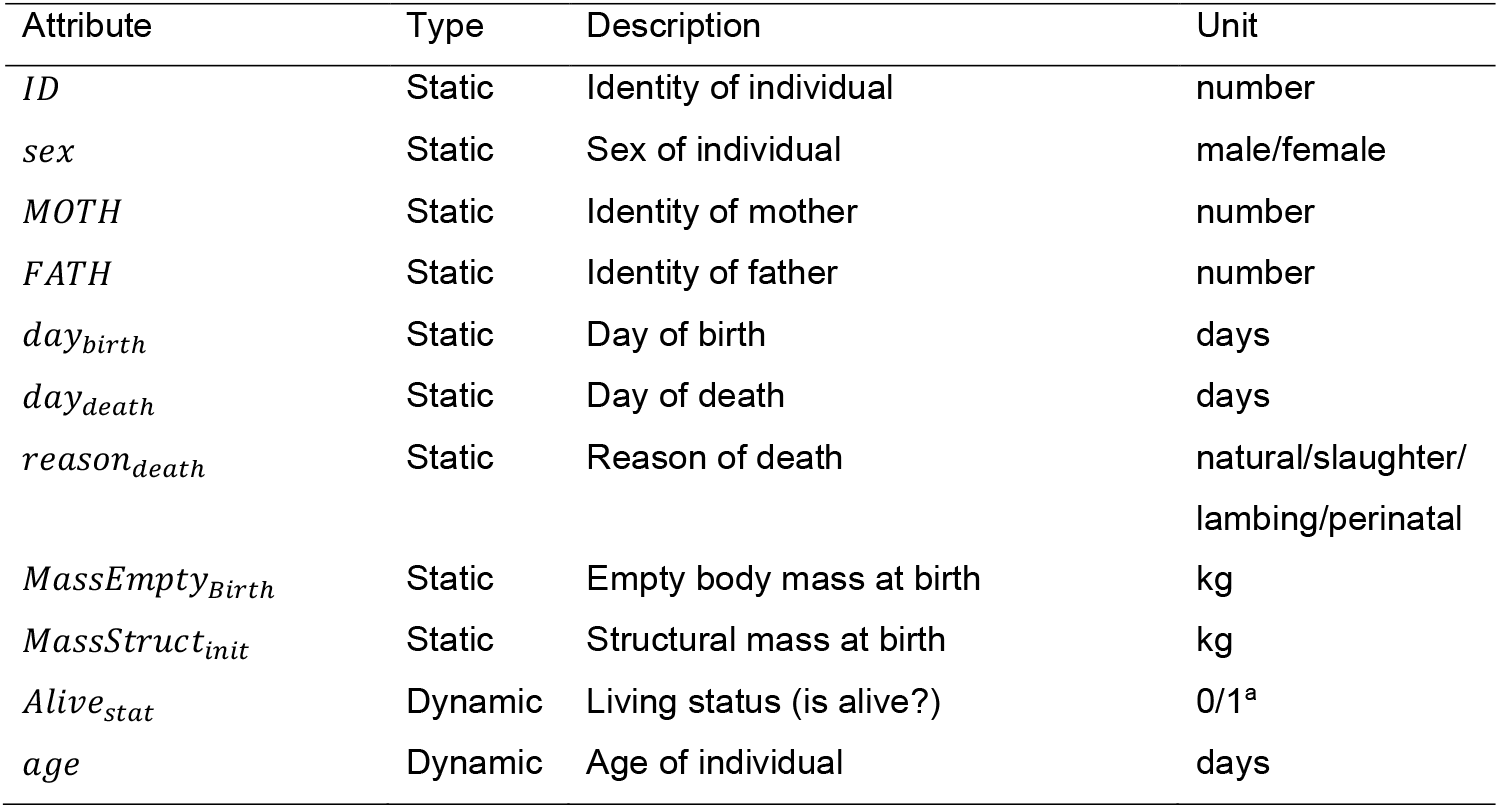
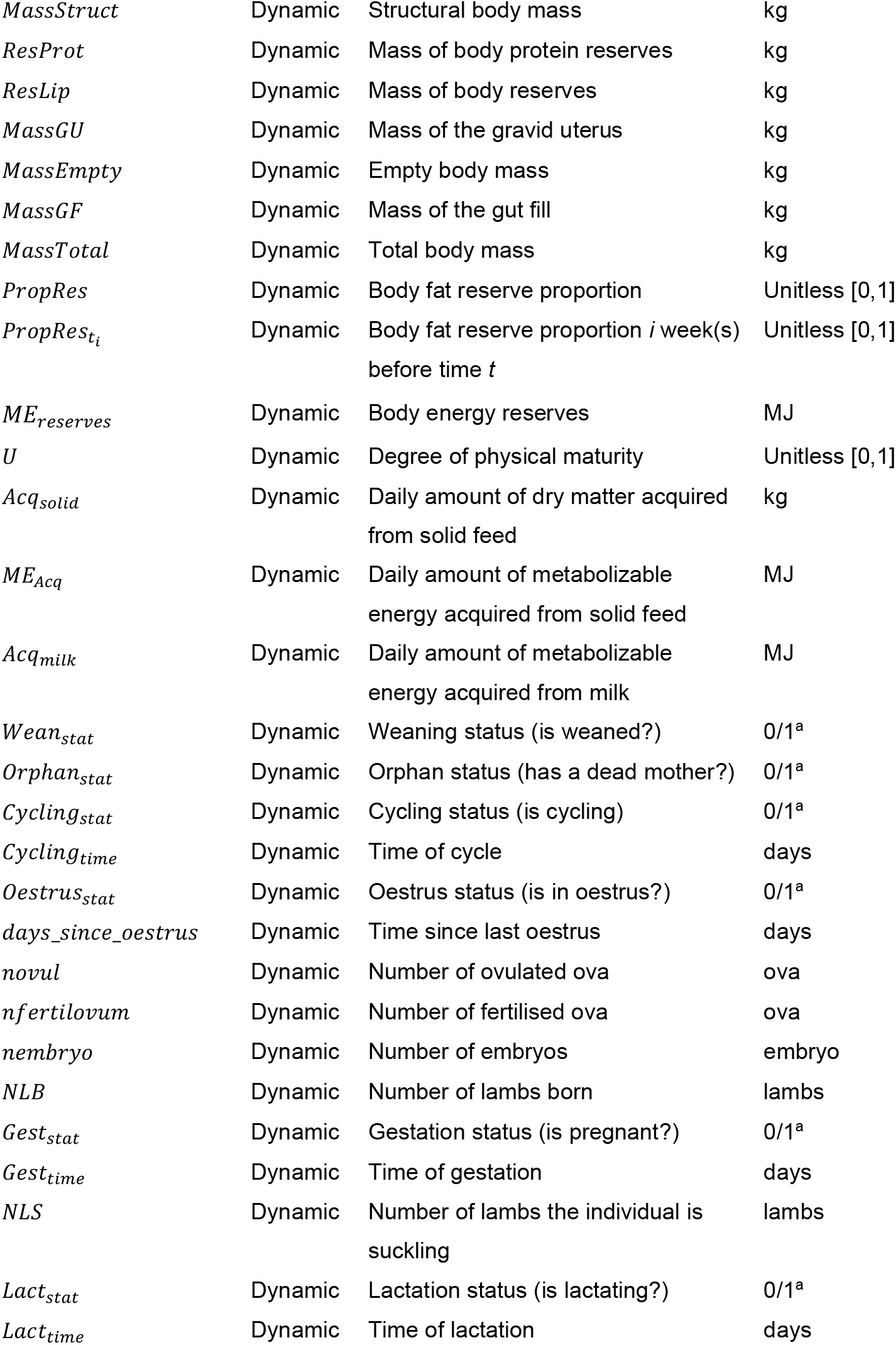
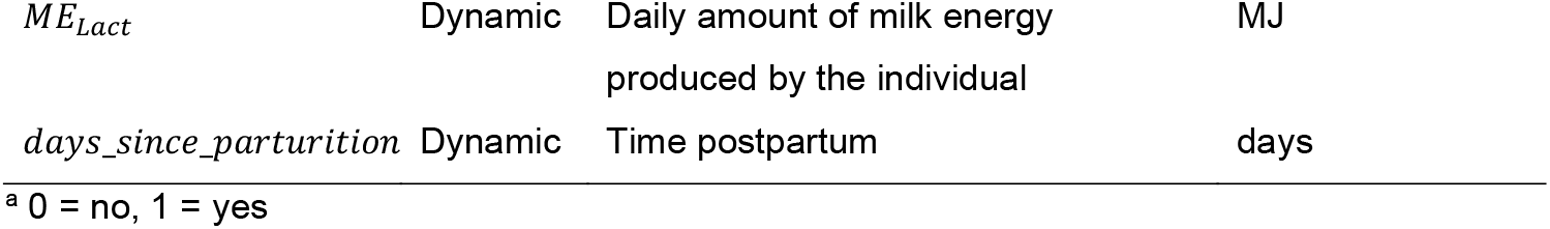
Definition of dynamic and static attributes of the sheep as represented in the model.

The individual sheep is exposed to an environment defined by five components:

i. the feed energy content (*MEC* in MJ metabolizable energy / kg dry matter) and the available feed quantity (kg dry matter) for each day of the year (each year has 365 days, day 1 = first of January),
ii. the length of the day (in hours) for each day of the year (1 to 365) depending on latitude (−90° to 90°),
iii. the reproductive management defined by the days of the year where a male is present and the minimal age at mating for ewe lambs (in days),
iv. the lactation management defined by the length of the suckling period
v. the culling management defined by rules to remove ewes from the herd.

#### 3. Model process overview and scheduling

After initialisation, the model processes are executed at each day. First the environmental variables are updated from the desired inputs. Physiological status and times are updated. Weaning status of lambs is defined by age and prior to weaning, lambs are classified as suckling lambs if they can be associated with a lactating mother or as orphan if not. The different processes that make up the nutritional model (Acquisition, Allocation, Utilisation; Fig. 1) are then successively executed. Part of these processes are first executed for all individuals who are not suckling and then for suckling lambs. Specifically, the desired amount of energy acquired from solid feed or artificial milk in non-suckling individuals is calculated and more or less covered depending on the availability of these resources in the environment at that day. The energy acquired from feed is then allocated between biological functions according the sex and the physiological state of the individual or the litter size. Biological functions include somatic functions (maintenance and reserves), growth, pregnancy and lactation. For lactating ewes, the energy allocated to lactation determines a milk energy output that is allocated to their suckling lambs and split between siblings. Suckling lambs acquire energy from milk intake and possibly from solid feed if milk energy does not cover the desired amount and if the lamb has the capacity to eat solid feed. The total amount of energy that lambs acquire is allocated between somatic functions and growth. For all individuals (lambs and ewes), the process of energy utilisation is then executed. The energy requirements for maintenance are updated. The energy balance (*EB*) corresponds to the difference between energy intake and the energy used for maintenance, growth, pregnancy and lactation. It determines if energy is deposited to (*EB* > 0), or mobilised (*EB* < 0) from body energy reserves. Body components including the mass of body reserves, structure and the mass of the gravid uterus are updated based on the amounts of energy allocated and the efficiency of energy use. After nutritional processes, three events are evaluated: conception, birth and death. Conception and birth are executed by the reproduction sub-model. If the ewe is cycling and if oestrus occurs, the number of ovulations is sampled based on the ewe potential rate of ovulation and her body features. If at least one ovulation occurs and a male is present at this time of the year, fertilisation can occur. If at least one egg is fertilised and becomes a viable embryo, then pregnancy starts. A birth event occurs if an ewe is in her last day of pregnancy. In that case, a new individual corresponding to each foetus of the ewe lambing at this day is created. For each new individual, an *ID* is assigned and sex is randomly sampled. Individual birth weight is determined based on the mother’s mass of gravid uterus and litter size.

**Fig. 1:**
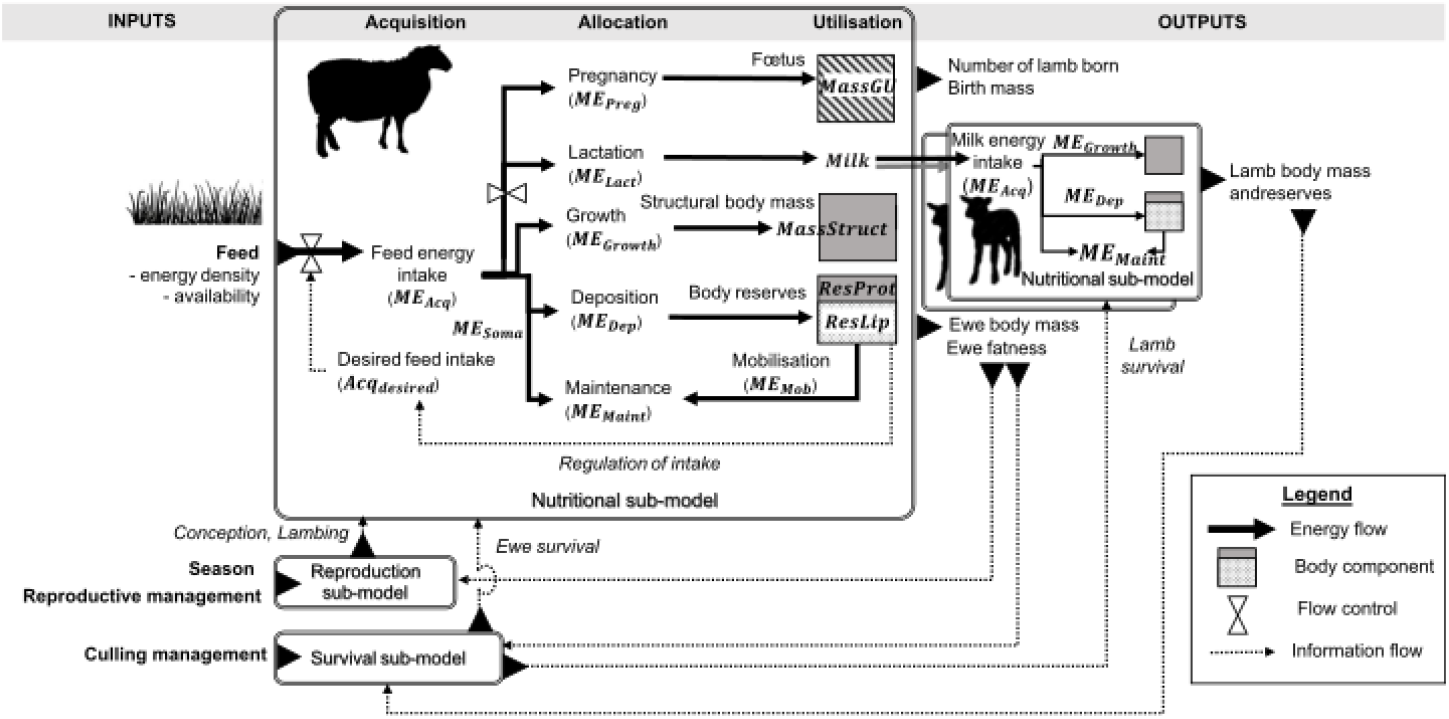
Conceptual diagram of the energy allocation model linking the ewe to her lambs. Energy intake acquired from feed is allocated between growth, pregnancy, lactation, maintenance and body reserves. Body reserves and reproductive status modulate energy acquisition and allocation. The use of energy allocated to growth and body reserves lead to changes in body tissues of the ewe while the energy allocated to pregnancy and lactation is transferred to her lamb(s).

In the survival sub-model, death events can occur at each day of an individual’s life. Newborn death is partly random as it is based on a survival function of birthweight. Death at any other age is also partly random but based on a general survival function of body reserves. Moreover, individuals can be removed according the culling criteria defined as input.

During the simulation from *t* = 0 to *t* = *TSIM*, all animals’ dynamic attributes are saved at each recording time-step *dT*.

#### 4. Design concepts

##### 4.1 Basic principles (concepts, theories, assumptions)

A central assumption of our model is that nutritional responses to the environment are based on the acquisition-allocation principle (Fig. 1). Following the law of conservation of energy, the amount of energy intake equals the amount of energy expenditure plus the variation in body energy reserves. Moreover, energy that is allocated to one biological function cannot be allocated to another function. Due to this allocation constraint, energy allocation trade-off may occur between functions. Importantly, in our model, maternal energy transfers are explicitly represented: energy allocated to pregnancy is transferred to lamb birth mass and energy allocated to lactation is transferred to suckling lambs with a potential competition among siblings if the total the total energy demand of the litter exceeds the amount of milk energy provided by the mother. Finally, responses can vary among individuals depending on a subset of user defined input parameters that directly or indirectly control energy acquisition and allocation processes. Among the different input parameters (Table 2), the subset of those varying among individuals is not identified a priori. We suggest that parameters associated with a target level (identified with a “*”) can underpin genetic variation in energy allocation (e.g. *AllocGrowth*^*^, *MassStruct*^*^, *AllocLact*^*^, *ovulrate*^*^). However other parameters may also be genetically variable and modulate energy allocation through time changes for instance (e.g. 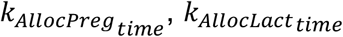).

**Table 2:**
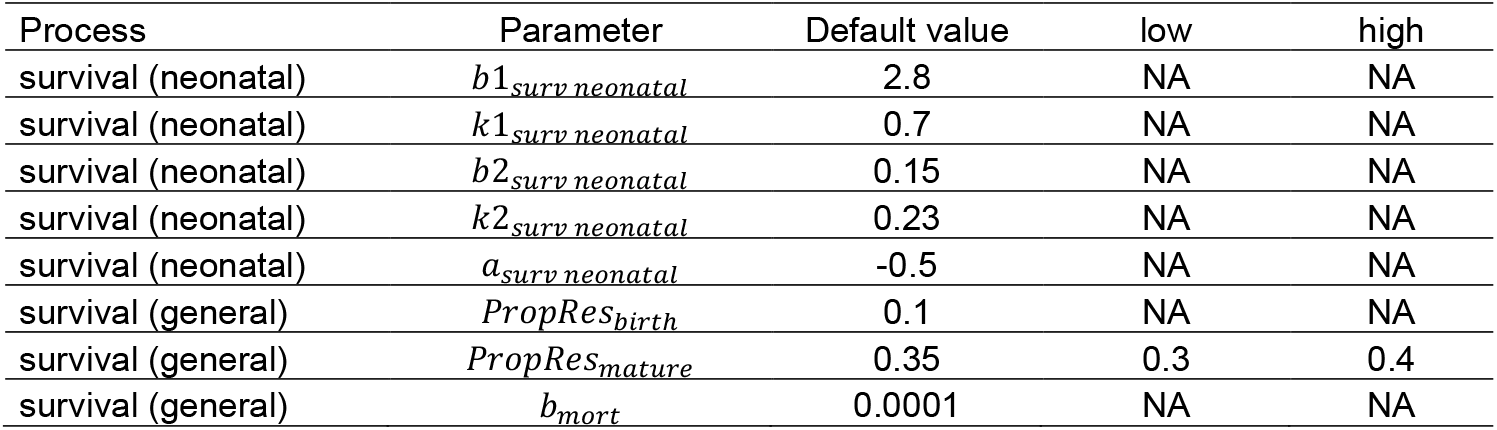
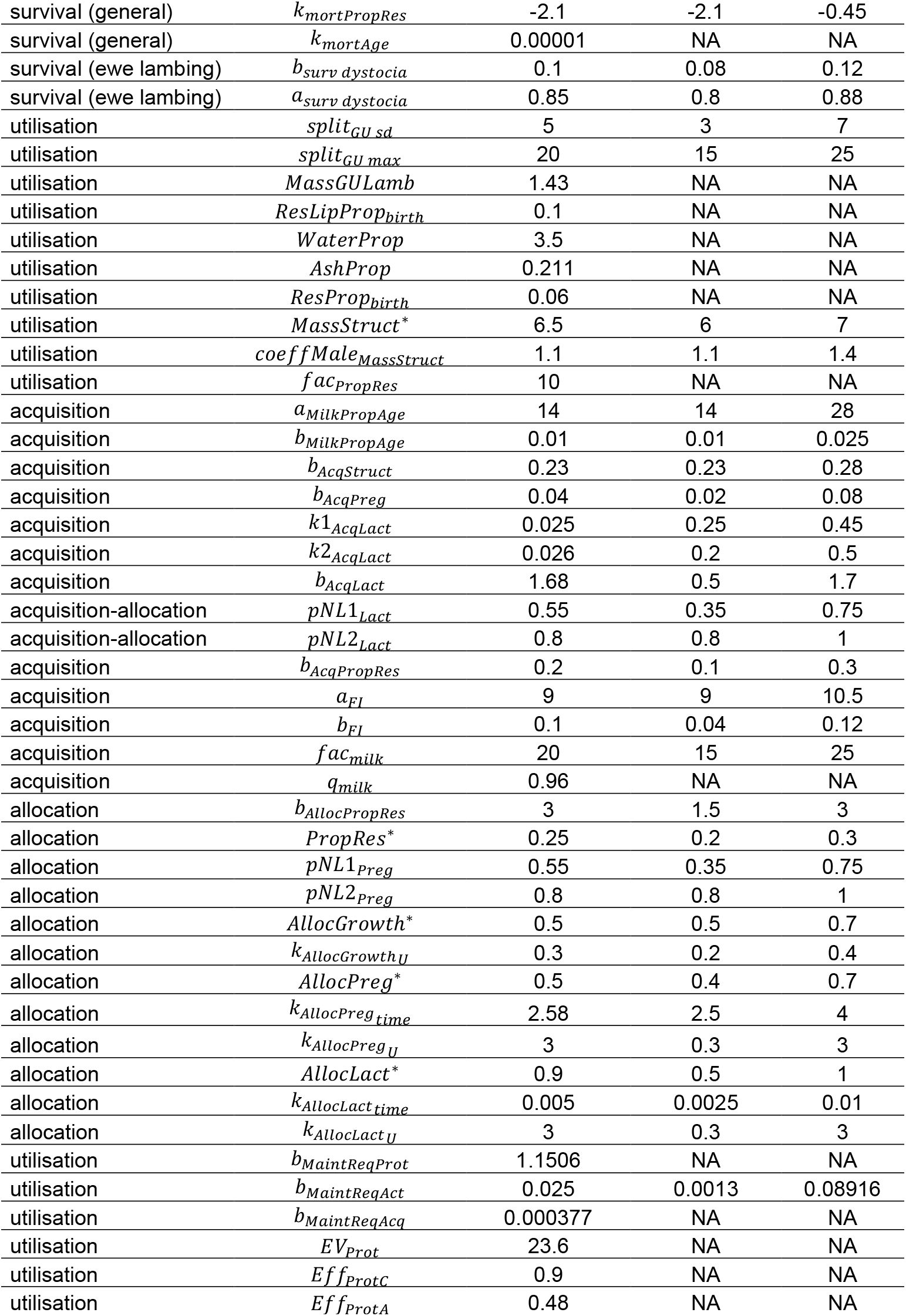
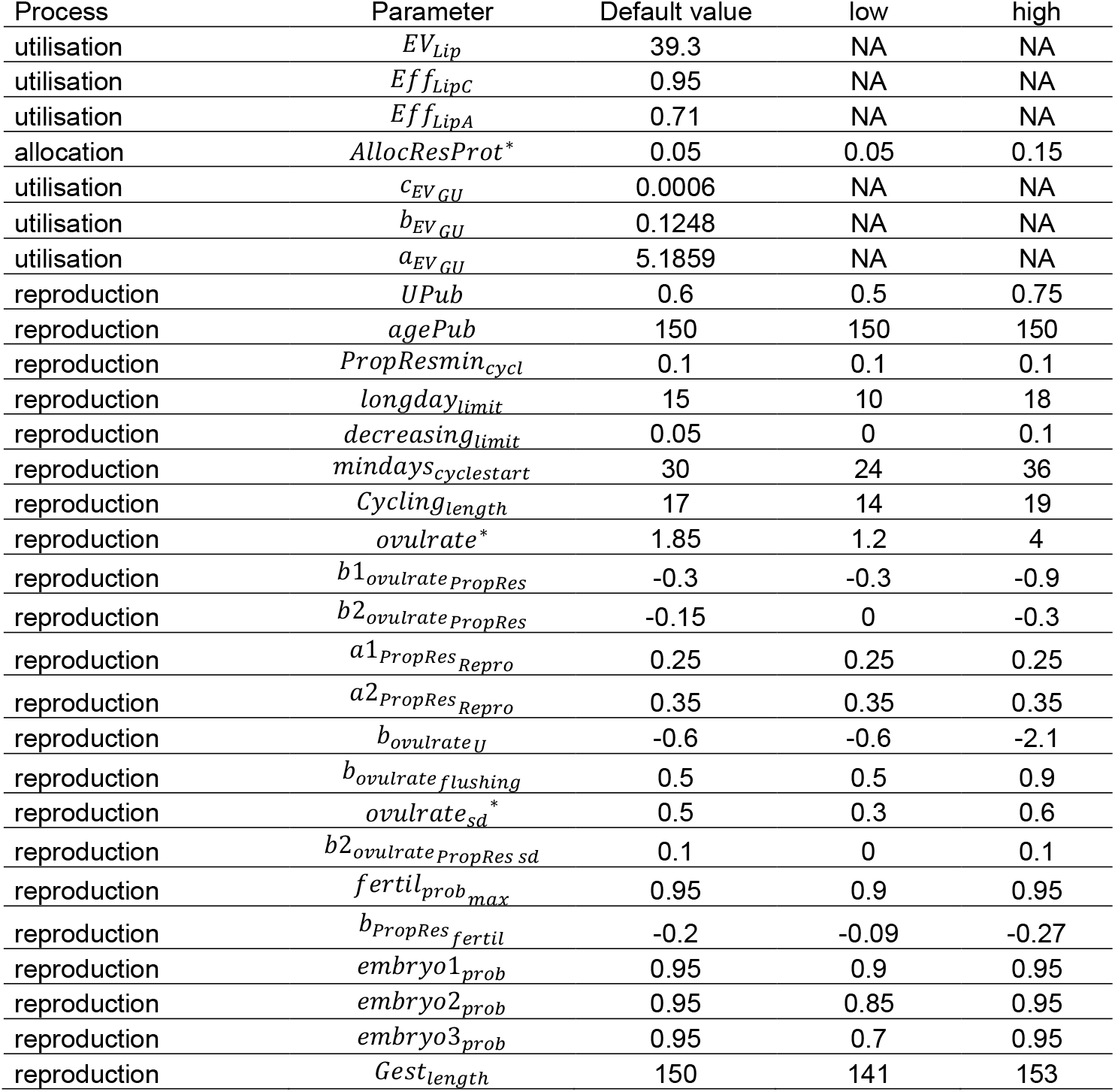
Sheep model input parameter default values and proposed ranges for variable input parameters.

##### 4.2 Stochasticity

Survival, reproduction and birth involve stochasticity to better take into account the random variation usually observed in those processes and that cannot be predicted deterministically here. For survival, individual attributes such as age and reserve proportion influence daily survival probabilities, lamb neonatal survival and ewe survival at birth due to dystocia. Reproduction encompasses a series of stochastic events including oestrus occurrence, number of ovulations, how many of those ova are fertilized and how many develop into embryos. At birth, the total mass of the litter is distributed among siblings using random sampling and each lamb’s sex is randomly sampled.

##### 4.3 Interaction

Interactions are only assumed between the ewe and her lamb(s) as well as among siblings before weaning. The ewe determines the lamb birth mass and the available milk. Reciprocally, the number of lambs shapes the ewe’s pregnancy and lactation by modulating the energy acquisition and allocation. Moreover, ewe’s survival probability at lambing is dependent on the lamb’s birth mass and of its survival.

Other interactions among individuals (e.g. competition for feed access) are currently not represented but they may be developed later.

##### 4.4 Emergence

Lifetime trajectories of the multiple traits emerge from the interaction between an individual and its environments, and from the interaction between the ewe and her lambs. Moreover, complex patterns may emerge at the level of a population or a flock made-up of various individuals (e.g. flock resilience) although this dimension is not explored in this study.

##### 4.5 Adaptation

Each day, each individual “desires” a particular amount of feed and then seeks to satisfy this desire. The processes of acquisition and allocation are adapted according to individual’s state, in particular its fat reserves proportion. The reproductive events that can lead to gestation are also adapted according to individual’s state.

##### 4.6 Objectives

In the adaptation process, the objective measure to evaluate alternative responses (e.g. reduce or increase energy acquisition) is the body fat reserve proportion (*PropRes*). When this proportion is below a target level (*PropRes*^*^), individuals increase their desired feed intake and vice versa reduce it when it is above. Similarly, when *PropRes* of a pregnant (or lactating) ewe is below *PropRes*^*^, then energy allocation to pregnancy (or lactation) is decreased to favour maternal survival at the expense of the offspring. The level of *PropRes* during the last six weeks before ovulation also influences the probability to start a gestation.

##### 4.7 Sensing

Each individual is assumed to sense the environmental conditions and the level of its reserves. Ewe sense the size of their litter to adjust their energy allocated to pregnancy or lactation. Additionally, lambs sense the amount of milk provided by their mother.

##### 4.8 Learning

In the ODD protocol, learning refers to individual that change how they produce adaptive behaviour over time as a consequence of their experience (Grimm et al., 2020). Currently, this concept is not implemented here.

##### 4.9 Prediction

We do not assume that individuals are able to explicitly predict future conditions to make their decisions. Implicitly though, the level of their body reserves (*PropRes*) and of its variation are assumed to inform individuals about future feed availability in their environment. In particular, this information partly determines the “decision” to start a gestation or not.

##### 4.10 Collectives

The model includes no collectives but can provide a basis for representing a flock or a population.

##### 4.11 Observation

Each individual’s static attributes (*ID, sex, MOTH, FATH, day*_*birth*_, *MassEmpty*_*Birth*_, *day*_*death*_, *reason*_*death*_) are recorded in a table describing all individuals from a simulation (initial animals and subsequent births). Individuals’ dynamic attributes (Table 1) at each user defined time-step (*dT*) are the longitudinal observations used to describe the trajectories of trait responses to feeding and management. Output traits trajectories can be used to characterize the individual’s allocation patterns defined by a combination of input parameters. However, the simulation of replicates and the use of summary statistics is recommended to account for the variation in trajectories that can be caused by stochasticity.

#### 5. Initialisation

The user defines the five environmental components previously mentioned in Section 2 and the initial group of individuals. This includes the number of initial animals, their sex, day of birth, and birth mass. By default, the values of input parameters of lambs born during simulation are inherited from their mothers. Values of the input parameters characterizing each individual allocation can be kept constant among individuals to simulate replicates. Alternatively, various values of input parameters can be defined among initial individuals to simulate individual variation in model outputs. Input parameter default values and proposed ranges are given in Table 2. The initial animals are not subjected to the random sampling of survival at birth (neonatal survival).

#### 6. Input data

The feed environment data is an input file describing for each day (from *t* = 0 to *t* = *TSIM*) two variables: feed metabolizable energy content (*MEC*) in MJ metabolizable energy per kg feed dry matter and available feed quantity (*Food*_*available*_) in kg dry matter per animal). It can be thus be used to mimic seasonal variability within a year or interannual variability in feed quality. At a given day, values of *MEC* and *Food*_*available*_ represent the total diet (e.g. forage and concentrate) and are the same for all individuals although more specific scenario could be easily developed (e.g. specific diets for triplet-bearing ewes or for specific batches). Values for *MEC* can be found in feed databases. Assumptions of the diet should be based on the production system that is represented. Vegetative growth models providing the amount of feed and its components to calculate the metabolizable energy content can also be used to generate the feed input.

#### 7. Sub-models

The individual sheep model is made up of three sub-models (Fig. 1): (1) the nutritional sub-model at the heart of the whole model and which contains the 3 processes of energy acquisition, allocation and utilisation, and (2) the reproduction sub-model and (3) the survival sub-model that predict the occurrence of reproductive and mortality events, respectively.

##### 7.1 Nutritional sub-model

###### 7.1.1 Acquisition

The main output of the acquisition process is the amount of energy *ME*_*Acq*_ that is subsequently allocated between functions. We first define *ME*_*Acq*_ for adult individuals and then for unweaned lambs.

After weaning, all energy intake is acquired from an amount of solid feed *Acq*_*solid*_ of energy content *MEC* (in MJ/kg dry matter):

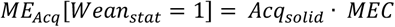

*Acquisition and feed quality:* In addition to the limitation due to *MEC*, low feed digestibility also reduces *Acq*_*solid*_ because of lower palatability of the feed and increased retention time (Pulina et al., 2013; Sauvant et al., 1996). As digestibility is correlated with feed energy density (*MEC*), we assume that *Acq*_*solid*_ is reduced at lower feed energy content (*DigestConstraint*). Additionally, actual feed intake is possibly limited by the amount of feed available (*Food*_*available*_). Actual solid feed intake (*Acq*_*solid*_) is then defined as follows:

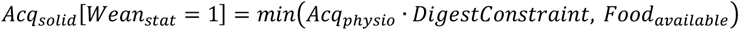

where intake capacity corresponds to the product between physiologically constrained acquisition (*Acq*_*physio*_) and *DigestConstraint* that is defined as follows:

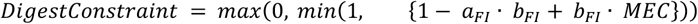

Except during pregnancy (see later), we assume that *Acq*_*physio*_ is equal to the desired acquisition (*Acq*_*physio*_[*Gest*_*stat*_ = 0] = *Acq*_*desired*_).

This demand *Acq*_*desired*_ increases linearly with *MassStruct* according to the factor *b*_*AcqStruct*_. Moreover we assume that body fatness acts as a negative feedback regulation of intake (Tolkamp et al., 2006). We formalize this effect as follows: when *PropRes* is higher than a target level *PropRes*^*^, then intake is reduced by the factor *b*_*AcqPropRes*_ and, if *PropRes* is lower than *PropRes*^*^, intake increases (Fig. 2). Desired feed intake for non-pregnant and non-lactating animals is thus defined as follows:

**Fig. 2:**
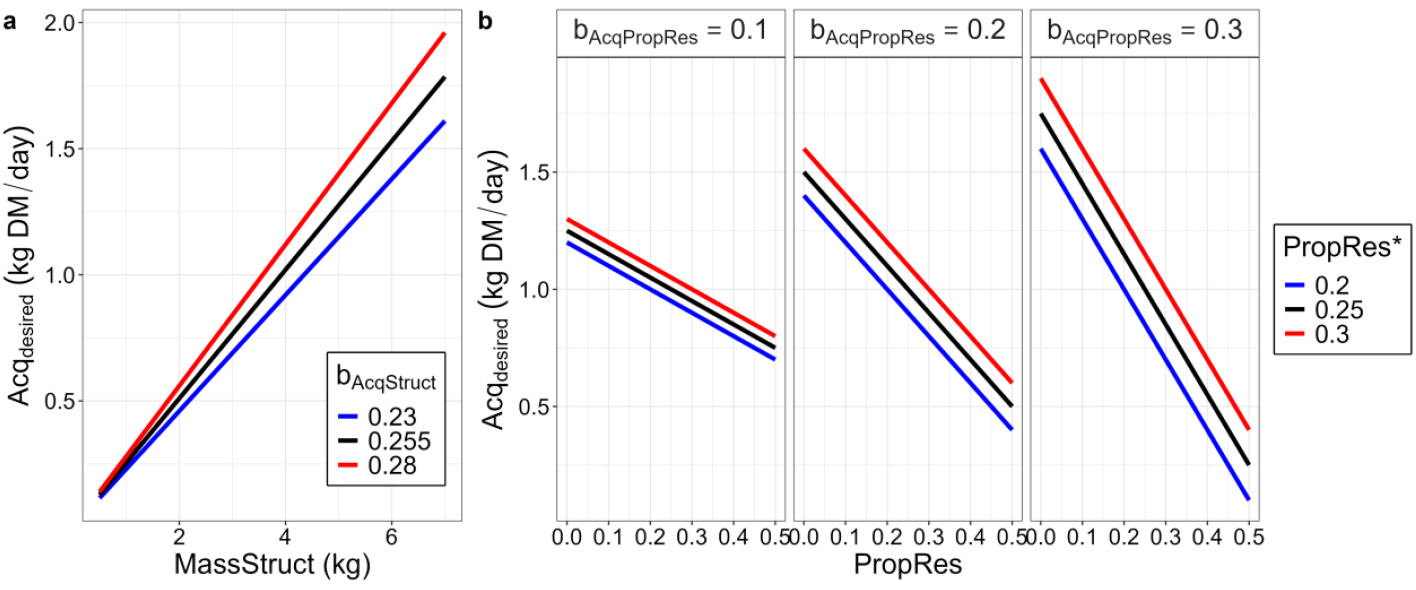
Effect of structural mass (*MassStruct*) and body fat reserves (*PropRes*) on desired feed acquisition (*Acq*_*desired*_). **a** *Acq*_*desired*_ in kg dry matter/day is proportional to structural mass *MassStruct* according to the coefficient *b*_*AcqStruct*_ (here *PropRes* = *PropRes*^*^). **b** a feedback of body reserves on intake is represented with a modulation of *Acq*_*desired*_ by the deviation of *PropRes* from the target level *PropRes*^*^ according to the coefficient *b*_*AcqPropRes*_ (here *b*_*AcqStruct*_ · *MassStruct* = 1).

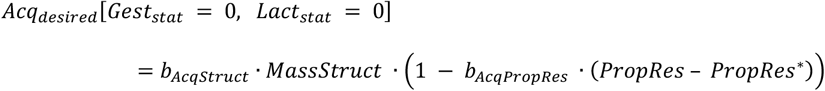

###### Desired feed intake increases during pregnancy and lactation

Pregnancy and lactation trigger an increase in the desired feed acquisition. In pregnant ewes, *Acq*_*desired*_ increases linearly with the gravid uterus mass *MassGU* according to the factor *b*_*AcqPreg*_. The increased intake during lactation is dependent on the number of lambs that the ewe is suckling (*NLS*) and the time of lactation (*Lact*_*time*_) (Peart, 1967). The general equation of *Acq*_*desired*_ including pregnant (*Gest*_*stat*_ = 1) or lactating ewes (*Lact*_*stat*_ = 1) is defined as follows:

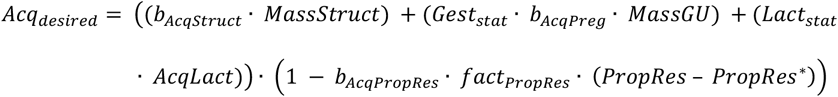

Additional intake *AcqLact* is assumed to change with lactation time *Lact*_*time*_ following a three compartmental model including two first-order reactions (compartment 1 → compartment 2 → compartment 3), where compartment 2 represents *AcqLact* dynamic (Fig. 3). The first compartment contains the initial amount *b*_*AcqLact*_ that is transferred to compartment 2 with fractional outflow rate *k*1_*AcqLact*_ before transfer to compartment 3 with fractional outflow rate *k*2_*AcqLact*_. The equation describing the dynamic of compartment 2 is further multiplied by the function *fprol* to modulate *AcqLact* according to suckling litter size (Fig. 4). If *k*1_*AcqLact*_ ≠ *k*2_*AcqLact*_ then *AcqLact* is then defined as follows:

**Fig. 3:**
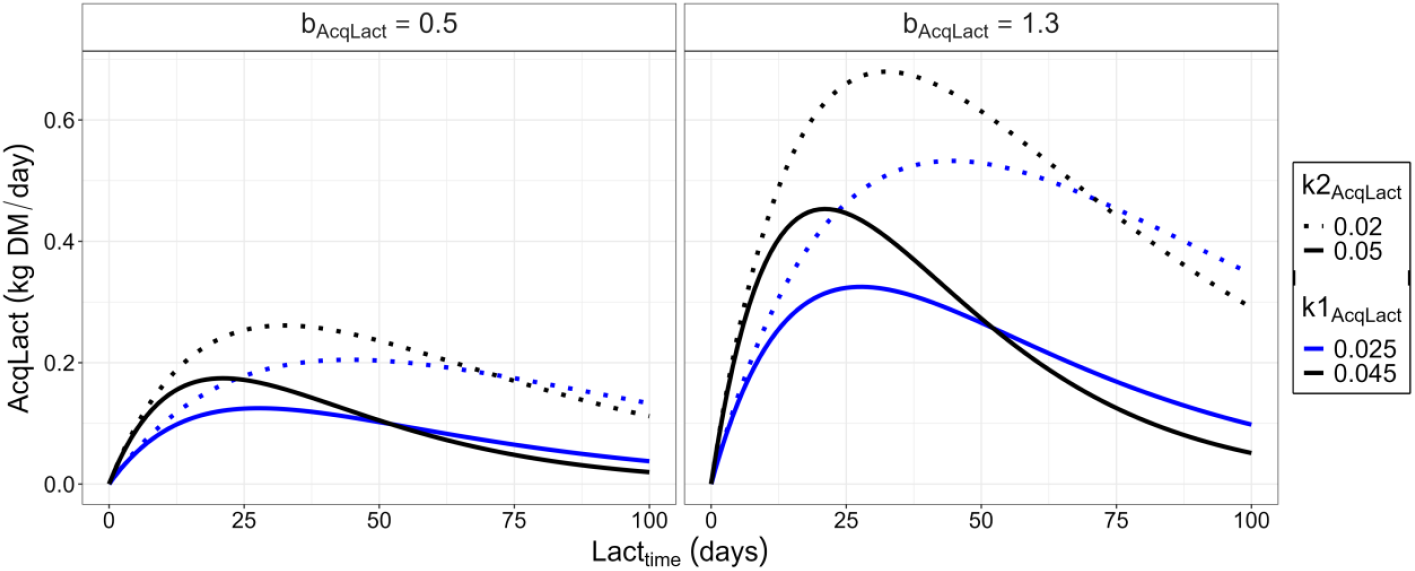
Effects of three parameters controlling the changes in additional desired feed acquisition (*AcqLact*) according to day of lactation (*Lact*_*time*_). Values are shown for an ewe suckling a single lamb with the coefficients *b*_*AcqLact*_ (left vs right panel), *k*1_*AcqLact*_ and *k*2_*AcqLact*_ (colour and line type). The maximum of *AcqLact* is reached when 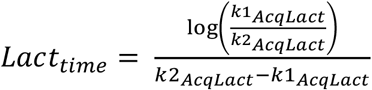.

**Fig. 4:**
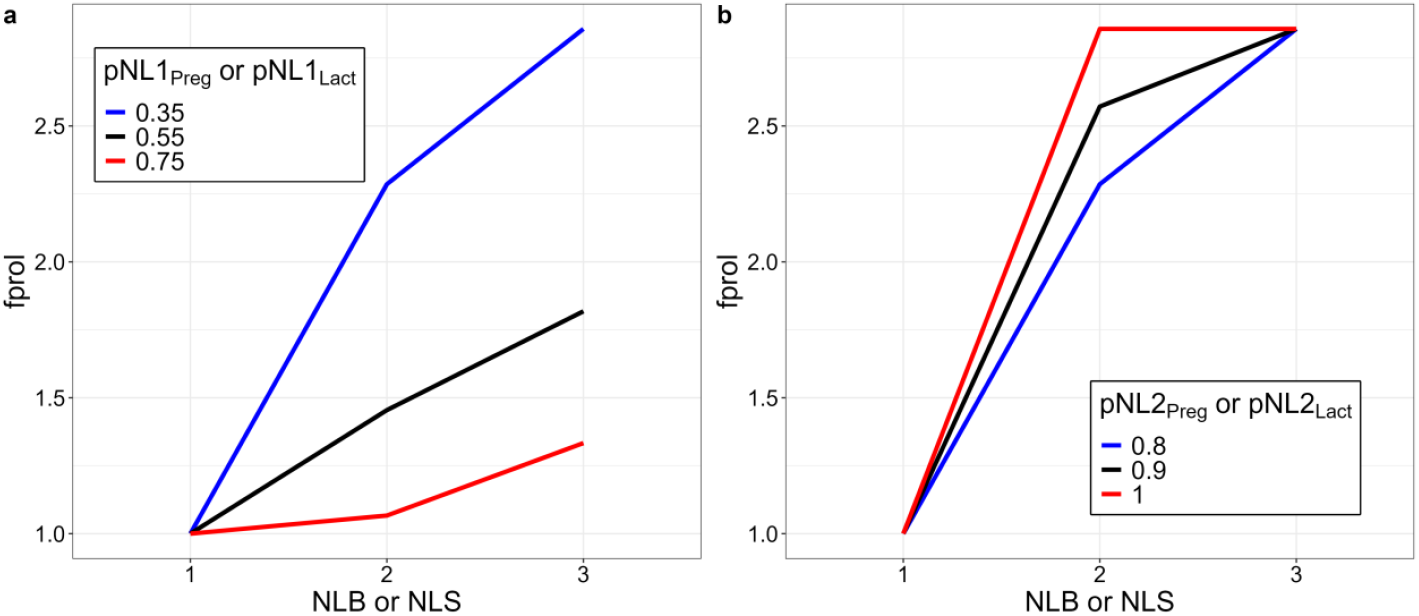
Modulation of acquisition and allocation (*fprol*) according to the number of lambs at birth (*NLB*) or suckling (*NLS*). The multiplier function *fprol* is controlled by parameters *pNL*1 and *pNL*2 **a** *fprol* decreases with increasing *pNL*1 (here *pNL*2 = 0.8). **b** The difference between 2 and 3 or more lambs decreases with increasing *pNL*2 (here *pNL*1 = 0.35).

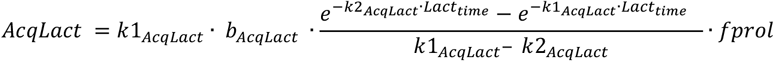

where

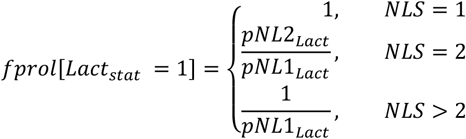

*Pregnancy constraint on feed acquisition:* The gravid uterus mass (*MassGU*) in relation to the structural body mass (*MassStruct*) and the empty body mass (*MassEmpty*) reduces intake at the end of gestation when the gravid uterus mass is high (Molina et al., 2001; Cooper et al., 1994; Wilkinson and Chestnutt, 1988; Forbes, 1968).This effect *PregConstraint* is stronger in ewes bearing multiple lambs (higher *MassGU*) or with high *MassEmpty* to *MassStruct* ratio (high body reserve proportion). Examples of *PregConstraint*, intake and body mass components are given in Fig. 5. Physiologically constrained intake (*Acq*_*physio*_) during pregnancy is defined as follows:

**Fig. 5:**
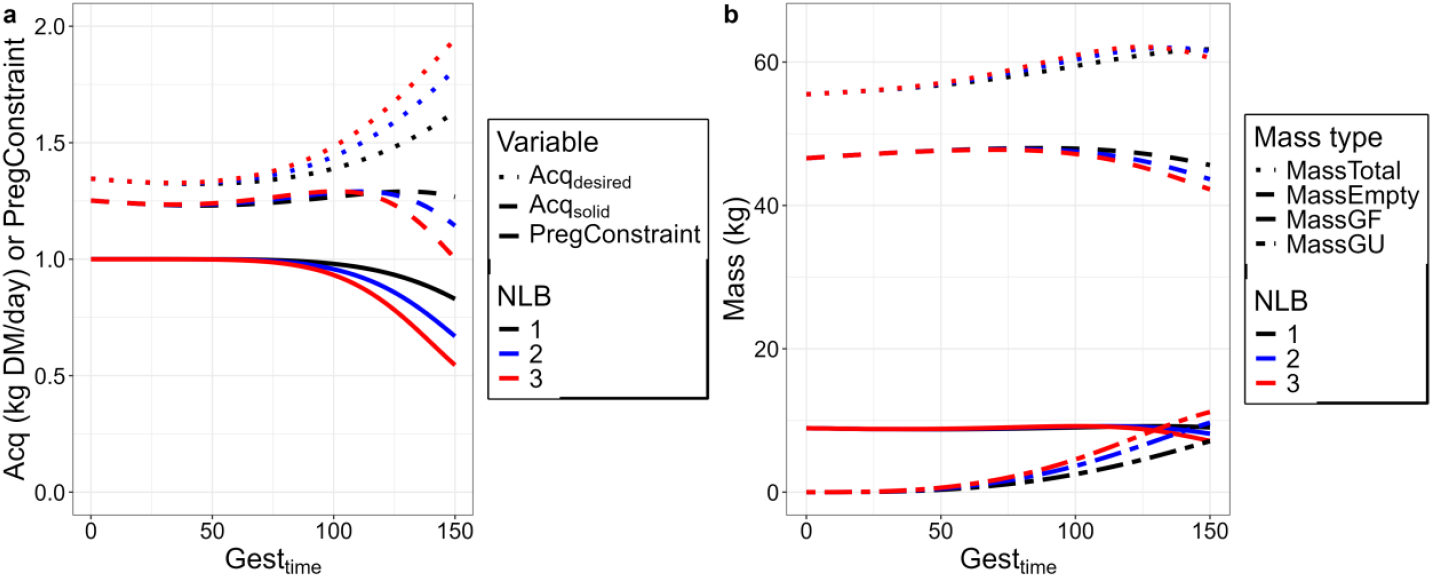
Effect of time of pregnancy (*Gest*_*time*_) and number of lambs born (*NLB*) on maternal feed acquisition (*Acq*) and components of body mass. **a** Due to the pregnancy constraint (*PregConstraint*), actual intake acquired from solid feed (*Acq*_*solid*_) is reduced at the end of pregnancy when *NLB* is high although desired acquisition (*Acq*_*desired*_) increases. **b** Total mass (*MassTotal*) increases during pregnancy due to the mass of gravid uterus (*MassGU*) whereas gut fill mass (*MassGF*) decreases because of reduced *Acq*_*solid*_ and empty body mass (*MassEmpty*) also decreases because body reserves are mobilized.

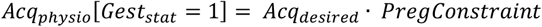

With

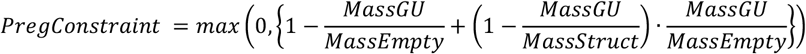

###### Solid feed and milk intake of the lambs

Before weaning, lambs acquire feed energy (*ME*_*Acq*_) from milk and from solid feed. Solid feed energy corresponds to a quantity of dry matter *Acq*_*solid*_ of energy content *MEC* while milk energy intake corresponds to *Acq*_*milk*_:

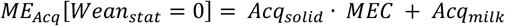

We assume that milk is prioritized over solid feed and that *Acq*_*milk*_ is driven by lamb’s desired feed intake. As for adults, desired feed intake in unweaned lambs depends on the structural body mass and the fat reserve proportion *PropRes*.

In suckling lambs, *Acq*_*milk*_ [*Wean*_*stat*_ = 0, *Orphan* = 0] is the minimum between the desired quantity 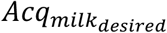 and the available quantity 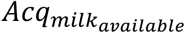 provided by its mother. The desired quantity of milk is expressed in energy with a factor *fac*_*milk*_ as follows:

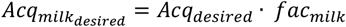

The available quantity depends on the mother’s allocation to lactation and the number of lambs in the litter. It is split between the siblings of the same litter (*NLS*) according to the proportion *propMilklitter*. Moreover, as the maternal milk energy (*ME*_*Lact*_) is a an amount of gross energy that is not fully metabolized by the lambs, it is then transformed into metabolizable energy with a conversion factor (*q*_*milk*_) (Jagusch and Mitchell, 1971). Therefore, milk energy available to the lamb of a litter is given by:

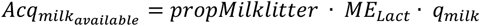

where *ME*_*Lact*_ is the maternal milk gross energy. The proportion attributed to sibling *I propMilklitter*[*sibling*_*i*_] is defined relatively to the total desired feed intake of the litter as follows:

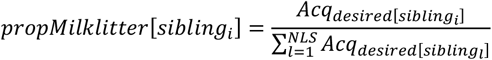

In orphan lambs, such as the initial lambs, artificial milk is assumed to be *ad libitum* and *Acq*_*milk*_ is assumed to change according to their intake capacity and the proportion of milk versus solid desired (*propMilkpred*).

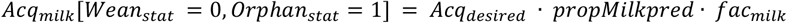

In all unweaned lambs, the proportion *propMilkpred* is assumed to linearly decrease with age (Doney and Peart, 1976), according to the coefficients *a*_*MilkPropAge*_ and *b*_*MilkPropAge*_ as follows:

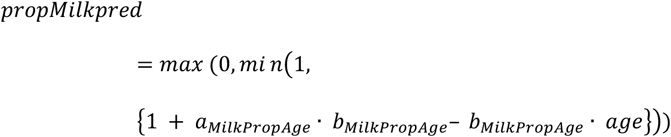

Solid feed intake is then defined as follows:

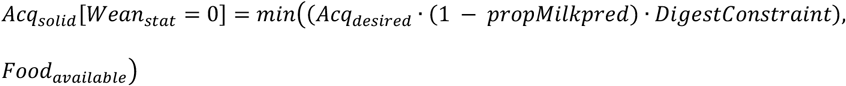

###### 7.1.2 Allocation

The acquired energy *ME*_*Acq*_ is possibly allocated to growth, pregnancy, lactation, maintenance and reserves deposition. The main outputs of the allocation process are the amounts allocated (*ME*_*Growth*_, *ME*_*Preg*_, *ME*_*Lact*_, *ME*_*Soma*_) that are available for the utilisation process. As a result of the allocation constraint, the sum of allocation coefficients (*AllocGrowth, AllocPreg, AllocLact, AllocSoma*) is equal to 1 and the value of each one is bounded between 0 and 1.

The feed energy allocated to each body function is the product of the proportion allocated to each function and *ME*_*Acq*_:

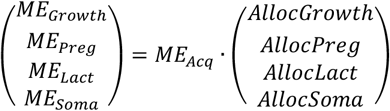

*Allocation to growth:* Allocation to growth (*AllocGrowth*) is determined by the basal allocation rate to growth (*AllocGrowth*^*^) and exponentially decreases with maturity *U* (Fig. 6). Physical maturity *U* is defined as the ratio between structural mass (*MassStruct*) and a target structural mass as fully grown adult (*MassStruct*^*^), both adjusted by structural mass at birth (*MassStruct*_*init*_). The allocation coefficient *AllocGrowth* is defined as follows:

**Fig. 6:**
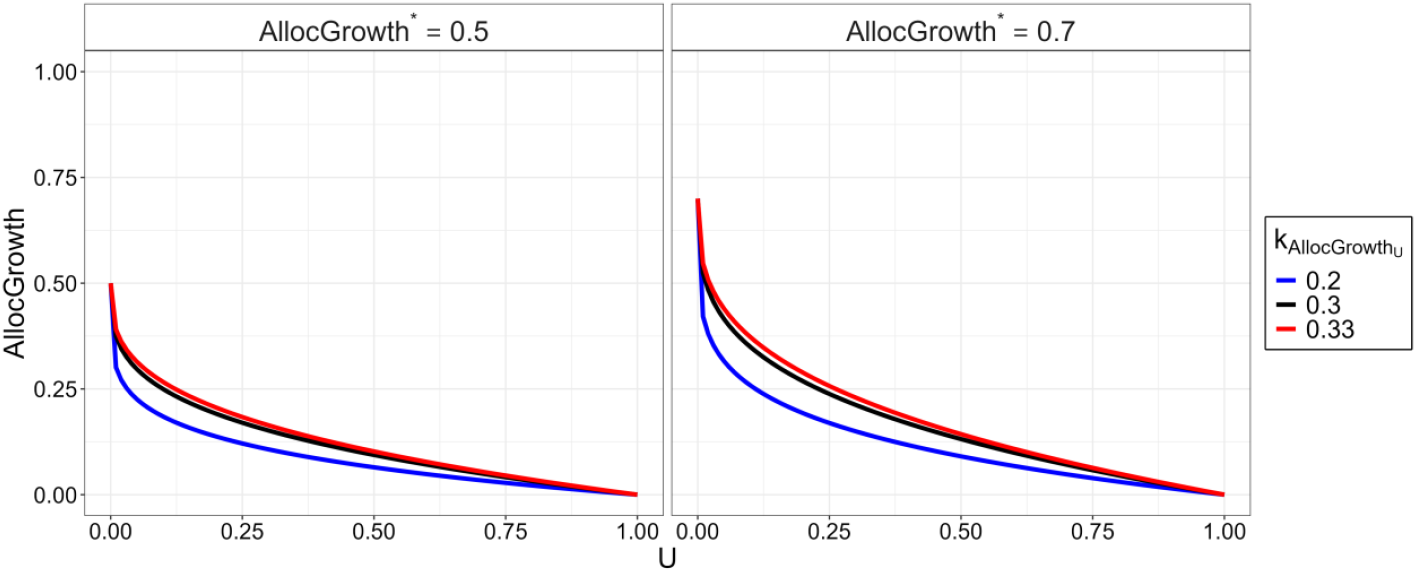
Effect of physical maturity (*U*) on allocation to growth (*AllocGrowth*). Allocation to growth *AllocGrowth* has a maximum *AllocGrowth*^*^ when *U* = 0 and exponentially decreases with *U* according to rate 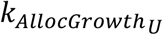.

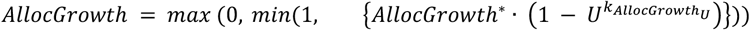

With

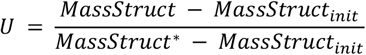

Sexual dimorphism in body mass is assumed through the factor *coeffMale*_*MassStruct*_ applied to *MassStruct*^*^ as follows:

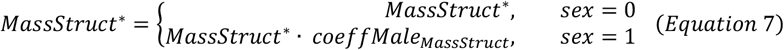

###### Allocation to pregnancy

Allocation to pregnancy (*AllocPreg*) is determined by a basal allocation rate to pregnancy (*AllocPreg*^*^), litter size during pregnancy (*NLB*), days of gestation (*Gest*_*time*_), maturity (*U*), and *PropRes*. It is defined as follows:

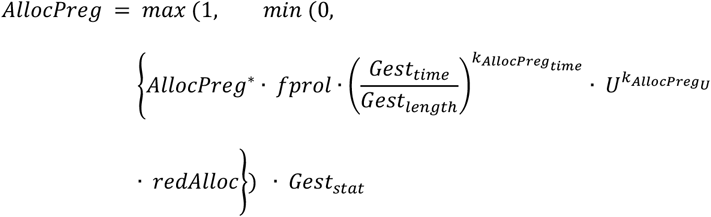

Basically, *AllocPreg* exponentially increases with *Gest*_*time*_ at a rate 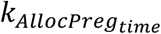 and up to the target level *AllocPreg*^*^ (Fig. 7). The target level is modulated according to litter size at birth (*NLB*) with the function *fprol* as follows:

**Fig. 7:**
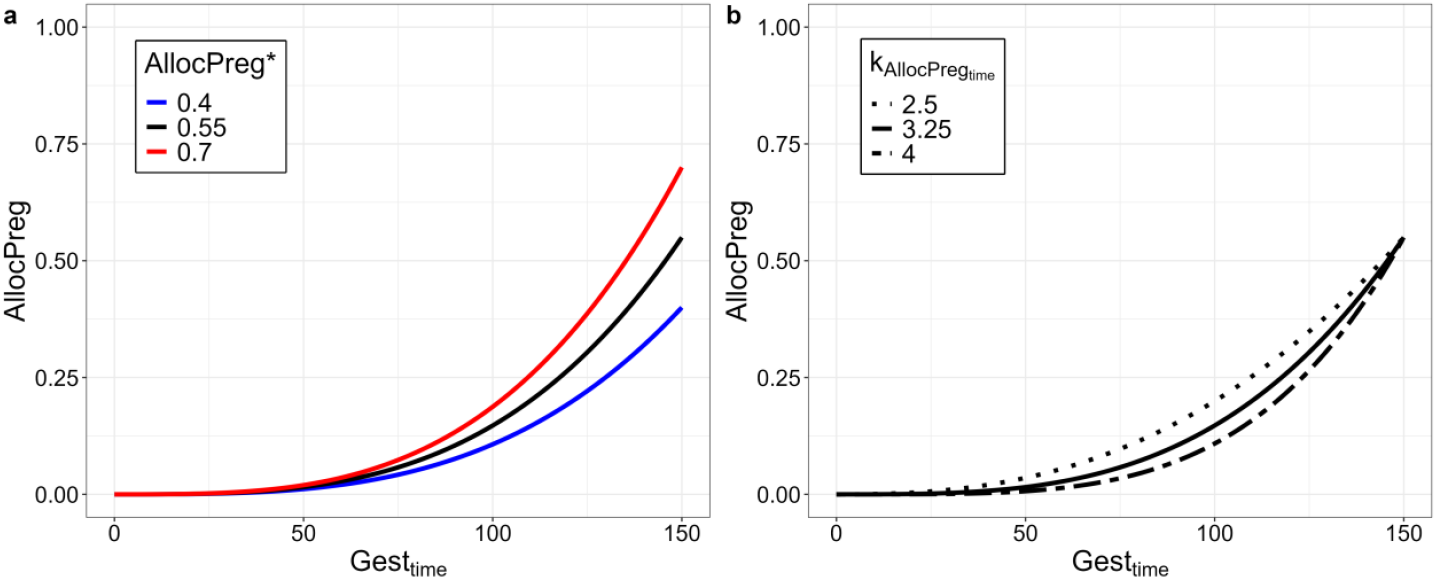
Effect of time of gestation (*Gest*_*time*_) on allocation to pregnancy (*AllocPreg*). *AllocPreg* is shown for *NLB* = 1, *U* = 1, *PropRes* = *PropRes*^*^. **a** Maximum allocation to pregnancy is determined by *AllocPreg*^*^ (here exponential decay 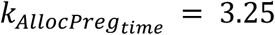). **b** The shape of the curve is determined by 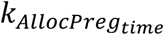 (here *AllocPreg* 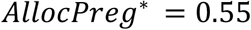).

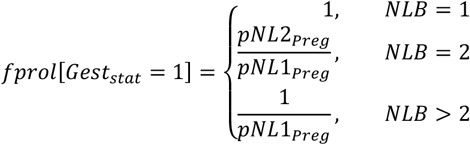

Allocation to pregnancy may also be reduced in young growing females. This effect is represented by the modulation 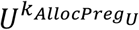 in the equation defining *AllocPreg*. Finally, when maternal reserves start being low, the priority of allocation to pregnancy is reduced. This reduction *redAlloc* is represented by a logistic decrease in *AllocPreg* when *PropRes* becomes low compared to *PropRes*^*^ (Fig. 8) as follows:

**Fig. 8:**
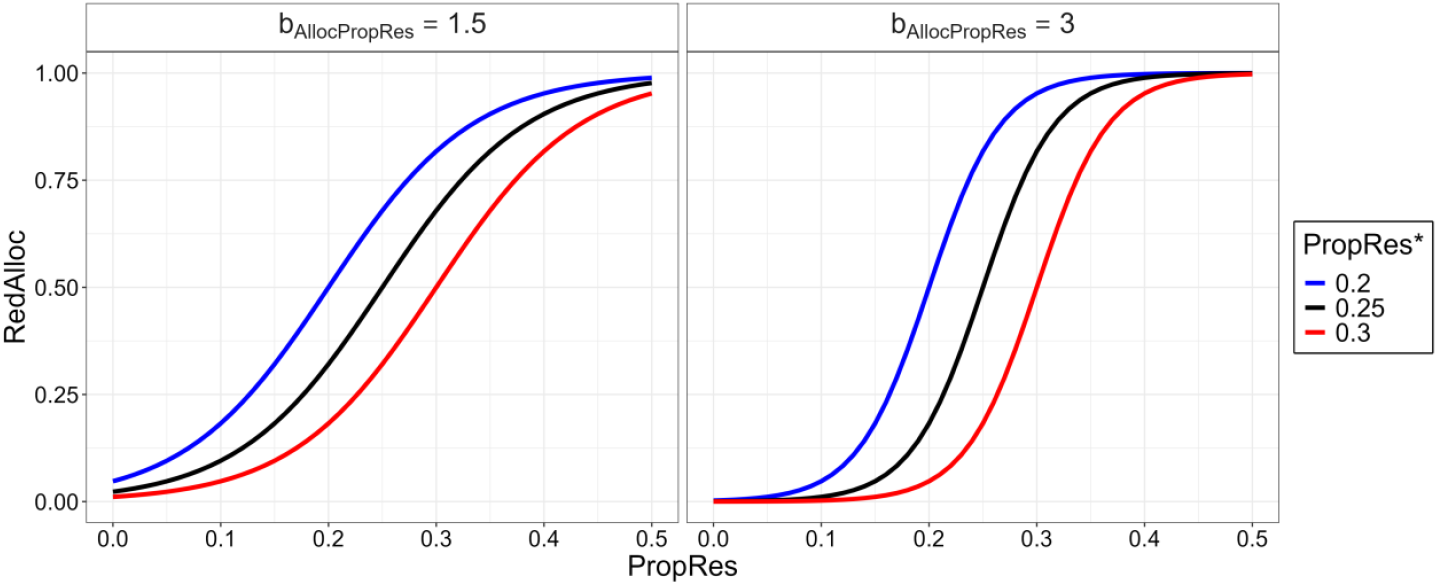
Effect of body fat reserves (*PropRes*) on reduction function of allocation to pregnancy or lactation (*redAlloc*). Allocations are reduced with the slope *b*_*AllocPropRes*_ when *PropRes* is below the target level *PropRes*^*^.

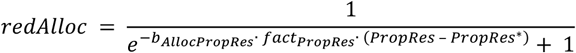

###### Allocation to lactation

Allocation to lactation (*AllocLact*) is determined by the basal allocation rate to lactation (*AllocLact*^*^), suckling litter size (via the function *fprol*), the stage of lactation (*Lact*_*time*_), maturity (*U*), and *PropRes*.

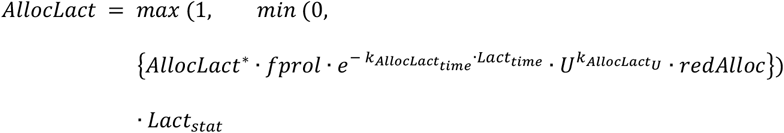

Basically, *AllocLact* exponentially decreases with time of lactation at a rate 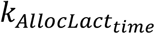 starting from the target level *AllocLact*^*^ (Fig. 9). This level is modulated according to suckling litter size (*NLS*) with the function *fprol* as previously shown. Finally, as for pregnancy, *AllocLact* is possibly reduced when body reserves drop (with the factor *redAlloc* as for *AllocPreg*).

**Fig. 9:**
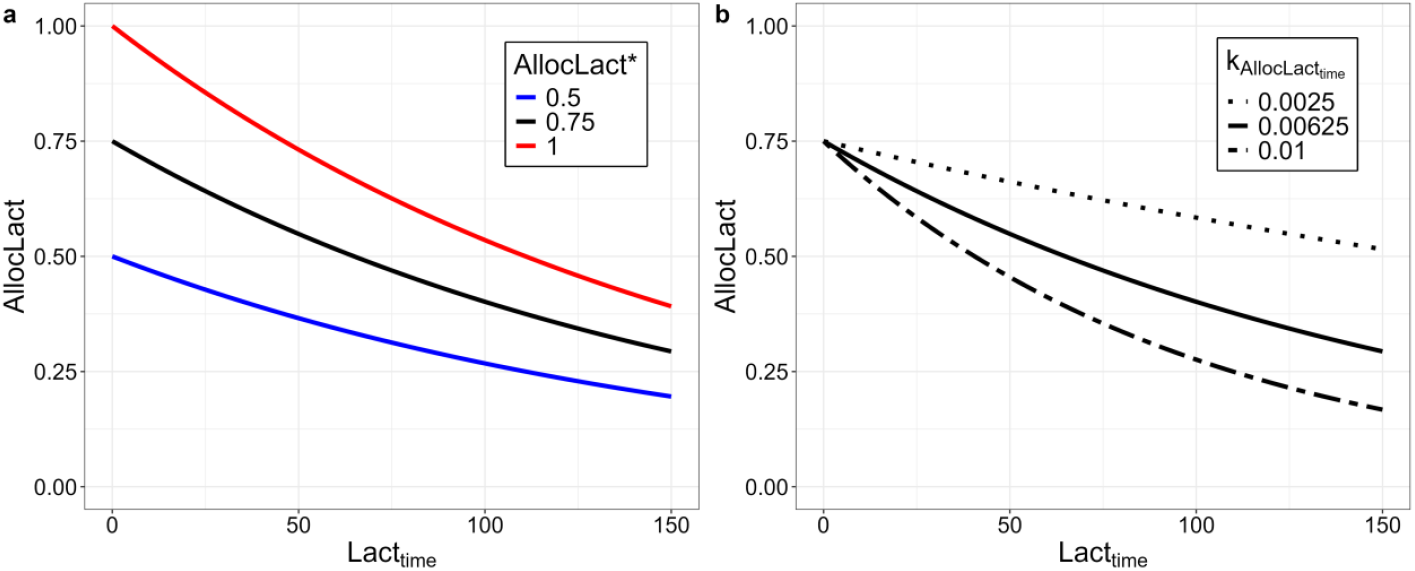
Effect of days of lactation (*Lact*_*time*_) on allocation to lactation (*AllocLact*). *AllocLact* is shown for a suckling litter size of 1, *U* = 1 and *PropRes* = *PropRes*^*^. **A** Maximum allocation to lactation is determined by *AllocLact*^*^ (here exponential decay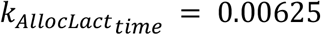). **b** The shape of the curve is determined by 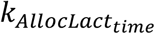 (here *AllocLact*^*^ = 0.75).

###### Allocation to somatic functions

Allocation to somatic functions (*AllocSoma*) includes self-maintenance and reserves deposition. It is determined according to the previous allocations *AllocGrowth, AllocPreg* and *AllocLact* as follows:

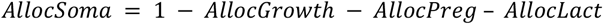

Note that when allocations reproductive functions are reduced as a consequence of low body reserves (*redAlloc* < 1), this favours individual survival through an increase in *AllocSoma*.

###### 7.1.3 Utilisation

The amounts of energy allocated (*ME*_*Growth*_, *ME*_*Preg*_, *ME*_*Lact*_, *ME*_*Soma*_) previously defined are the inputs of the utilisation process. This process predicts the time changes in the different products: body lipid (*ResLip*) and protein (*ResProt*) reserves, structural mass (*MassStruct*), mass of the gravid uterus (*MassGU*) and milk. Another part of *ME*_*Soma*_ is used for maintenance and dissipated as heat.

###### Maintenance

Energy allocated to soma include functions of maintenance and reserves use:

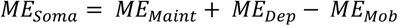

The variation in body energy reserves depends on the amount of *ME*_*Soma*_ relatively to the energy requirements for maintenance *MaintReq*. Once maintenance requirements *MaintReq* have been covered, the potential energy excess is allocated to reserves deposition (*ME*_*Dep*_ = *max*((*ME*_*Soma*_− *MaintReq*), 0)). In contrast, when *ME*_*Soma*_ is insufficient to cover maintenance requirements, energy is mobilized from the reserves if the amount of mobilizable energy reserves *ME*_*Reserves*_ is sufficient (*ME*_*Mob*_ = *min*((*ME*_*Soma*_− *MaintReq*), *ME*_*Reserves*_)). This situation of energy mobilization is likely to occur when *ME*_*Acq*_ is low or when a large proportion of it is used for pregnancy or lactation. At any time, the amount *ME*_*Reserves*_ is given by:

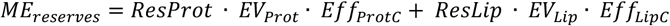

where *ResLip* and *ResProt* are body lipid and protein reserves, respectively, *Eff*_*Lipc*_ and *Eff*_*Protc*_ are catabolic efficiencies of lipid and protein reserves, respectively, and *EV*_*Lip*_and *EV*_*Prot*_are energy density of lipid and protein reserves, respectively.

In principle, *ME*_*Maint*_ = *MaintReq* if energy can be mobilized from *ME*_*reserves*_. However, when *ME*_*reserves*_drop then survival probability decreases towards 0. The energy requirement *MaintReq* is the sum of the needs for protein maintenance (*MaintReqProt*), the needs for activity (*MaintReqAct*) and the metabolic costs for energy acquisition (*MaintReqAcq*):

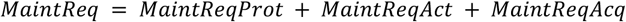

Those three types of requirements are proportional to protein mass, empty body mass, and energy intake, respectively:

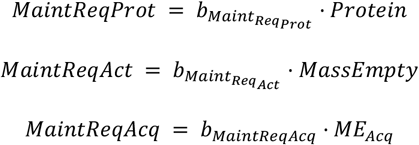

###### Reserves

The changes in the mass of protein reserves (*ResProt*) and lipid reserves (*ResLip*) between two consecutive days is given by (1) the storage of *ME*_*Dep*_to protein and lipid reserves according to the allocation coefficient *AllocResProt*^*^ and (2) the mobilization of *ME*_*Mob*_ from protein and lipid reserves according to the relative proportion *propLabProt*. This is defined as follows:

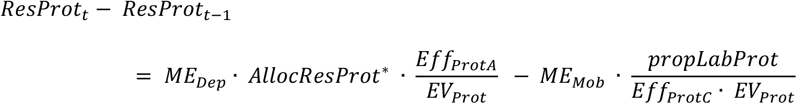

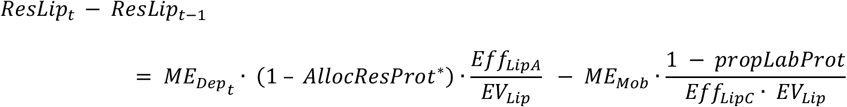

where

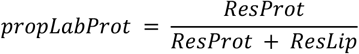

###### Growth

Growth in structural mass (*MassStruct*) is determined by the energy allocated to growth (*ME*_*Growth*_), the protein energy content (*EV*_*Prot*_), and the anabolic protein efficiency (*Eff*_*ProtA*_) as follows:

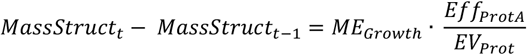

###### Gestation

Gravid uterus mass (*MassGU*) gain depends on the energy allocated to pregnancy (*ME*_*Preg*_) and the conversion efficiency (*EV*_*GU*_) which is dependent of the day of gestation (*Gest*_*time*_), as long as the ewe is pregnant (*Gest*_*stat*_ = 1):

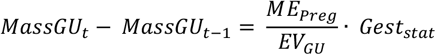

where:

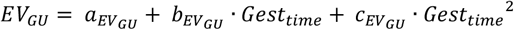

###### Milk produced

The metabolizable energy allocated to lactation (*ME*_*Lact*_) is directly equal the milk gross energy output of the ewe.

###### 7.1.4 Body components

Based on the updated mass calculated from the utilisation process, other body components are calculated.

The total body mass (*MassTotal*) is the sum of empty body mass, the gut fill and the mass of the gravid uterus in pregnant females:

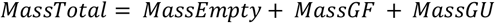

Empty body mass is further decomposed into its different chemical components as follows:

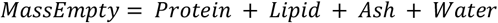

Protein mass includes structure and reserves (*Protein* = *MassStruct + ResProt*). Lipid is only considered as body reserves that can be stored or mobilized (*Lipid* = *ResLip*).

Ash mass is assumed to be proportional to the amount of protein constituting the body structure (*Ash* = *AshProp* · *MassStruct*).

Finally, water mass depends on the total protein mass (*Water* = *WaterProp* · *Protein*)

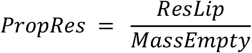

The body fat reserve proportion *PropRes* is the ratio lipid reserves to empty mass:

###### Gut fill mass

This mass (*MassGF)* is calculated from solid feed intake (*Acq*_*solid*_) and the metabolizable energy content (correlated to digestibility) of the feed (Laurenson et al., 2011), as follows:

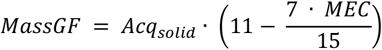

##### 7.2 Reproduction sub-model

###### 7.2.1 Cycling, oestrus and onset of pregnancy

The start of pregnancy occurs in a hierarchically organized sequence of events: cycling, oestrus, ovulation, fertilization, and embryonic survival lead to conception if at least one embryo is present after these events.

###### Cyclicity and reproductive season

Ewes need to fulfil conditions to be cycling. The animals need to be female (*sex* = 0). Minimum thresholds need to be reached for *PropRes* (*PropRes* ≥ *PropResmin*_*cycl*), maturity (*U* ≥ *UPub* maturity level needed for puberty), and age (*age* ≥ *agePub* minimal age needed for puberty) (Gómez-Brunet et al., 2011; Dýrmundsson, 1981). Pregnant (*Gest*_*stat*_ = 1) and lactating (*Lact*_*stat*_ = 1) ewes cannot be cycling. Ewes only resume cycling after a minimal recovery period after parturition (*mindays*_*cyclestart*_) (Gómez-Brunet et al., 2011).

The day of the year needs to be within the reproductive season determined by the ewe’s sensitivity to day length (determined by the day of the year and the latitude) and day length change (Robinson and Karsch, 1984; Hafez, 1952b, 1952a). Breeds differ in their sensitivity to day light (e.g. Forcada et al., 1992; Vosniakou et al., 1989; Wheeler and Land, 1977; Thimonier et al., 1969). The reproductive season is defined as follows (Fig. 10):

**Fig. 10:**
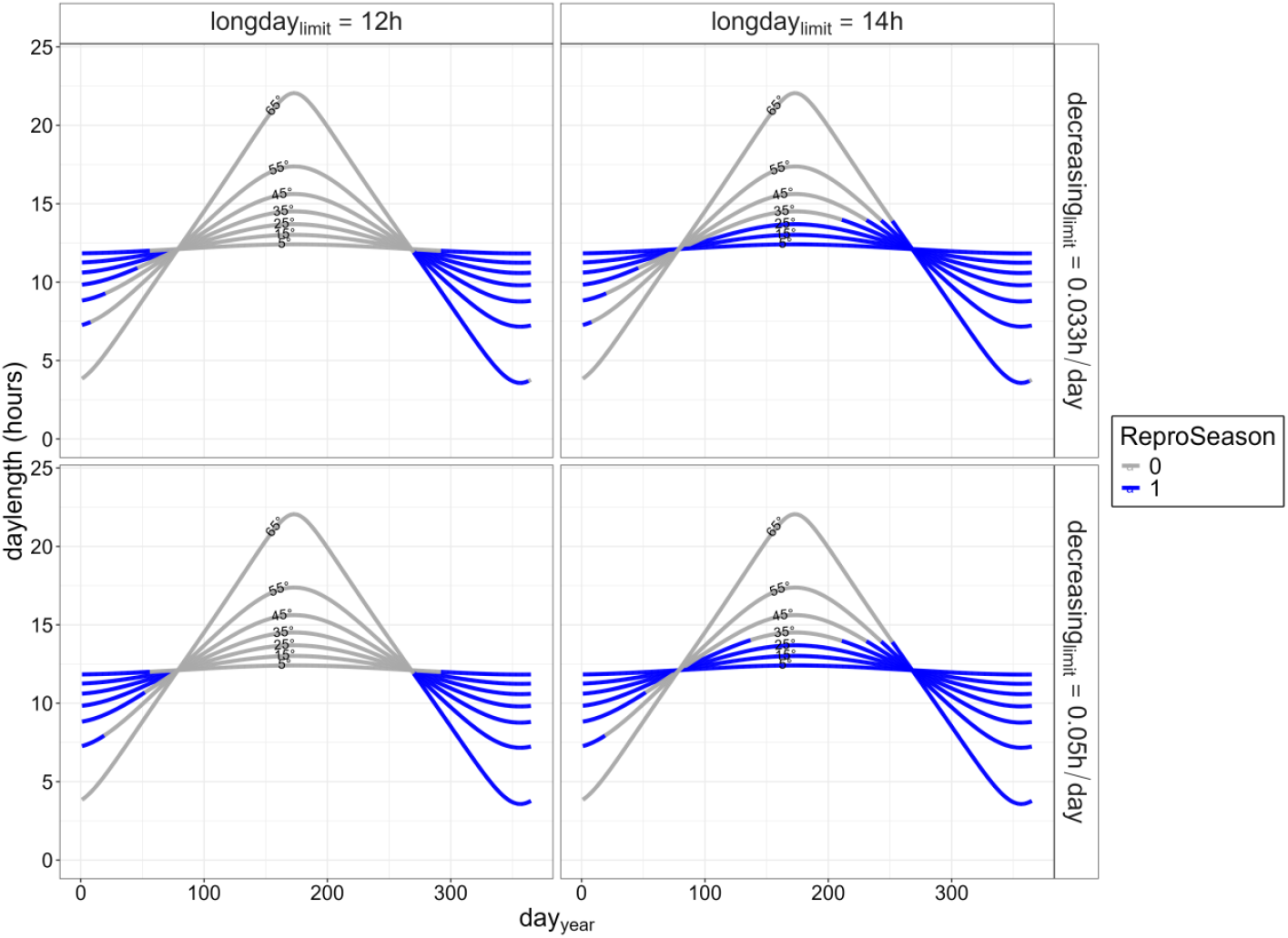
Reproductive season (*ReproSeason*) according to day of the year (*day*_*year*_) and latitude for different seasonality parameters. The day of the year and the latitude (in degree north) determines the length of the day (*daylength*). The hours of daylight and their change determines *ReproSeason*. Differences in seasonality are shown by varying the limit day length (*longday*_*limit*_) and the day length change limit (*decreasing*_*limit*_). The *refraction*_*period*_ was fixed at 30 days.

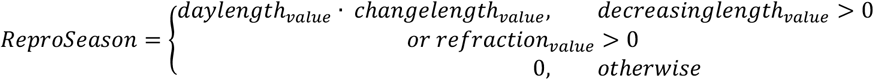

with:

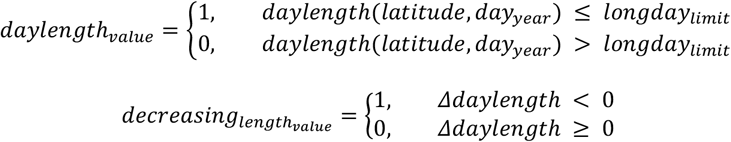

where *Δdaylength* = *daylength*(*latitude, day*_*year*_) − *daylength*(*latitude, day*_*year*_ − 1)

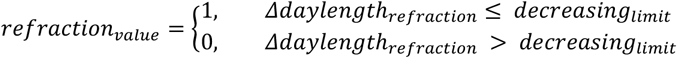

where

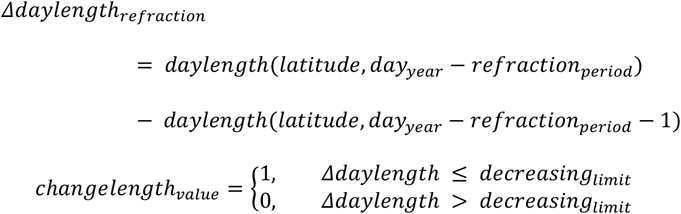

The latitude is the latitude in degrees (0 = equator, 0 < latitude ≤ 90: northern hemisphere, −90 ≤ latitude < 0: southern hemisphere), *day*_*year*_ the day of the year, *longday*_*limit*_ the day length maximum (hours) before resumption of cycling activity after a period of long days, the *decreasing*_*limit*_ the day length change limit below which the cycling activity stops (hours/day), and the *refraction*_*period*_ the number of days of refraction period in the period where the daylength increases.

###### Oestrus

Once the ewe has started cycling (*Cycling*_*stat*_ is changed from 0 to 1), each day, sampling from a binomial distribution determines if the first oestrus occurs or not. Oestrus probability increases with the number of days since the ewe started to be cycling (*Cycling*_*time*_) until reaching 1 at the duration *Cycling*_*length*_:

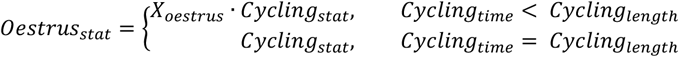

where 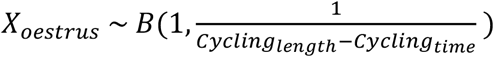

The day after oestrus, *Oestrus*_*stat*_ is set to 0. After each oestrus, the days since oestrus (*days*_*since oestrus*_) are counted. Subsequent oestrus happens after one cycle length if the ewe is cycling:

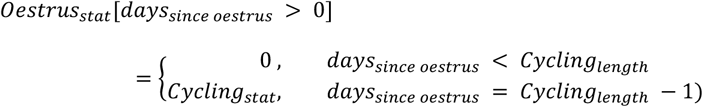

###### Ovulation

On the day when oestrus occurs, the number of ova novul is a positive integer number rounded after random sampling from a normal distribution of mean ovulrate_expect and SD sd_ovulrate:

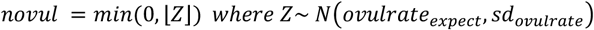

The expected ovulation rate ovulrate_expect_ depends on a target ovulation rate (*ovulrate*^*^). This expected ovulation rate is genetically determined, representing breed or genotype differences (Gootwine, 2020; Vinet et al., 2012; Davis, 2005). This rate is modulated by the body fat reserve level (Forcada et al., 1992; Gunn and Doney, 1975, 1979; Gunn et al., 1969). It decreases if *PropRes* are below the lower limit (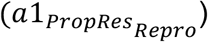) or above the upper limit (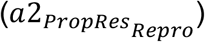) by the factors 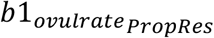 and 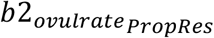,respectively (Fig. 11a). A gain in body condition (e.g. following flushing) can increase the ovulation rate in thin ewes (Rhind et al., 1984; Allison, 1975; Allen and Lamming, 1961). If *PropRes* is below the lower limit, the expected ovulation rate is changed by the factor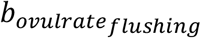 times the mean *PropRes* change 6 weeks to 1 week before the oestrus (*deltaPropRes*). Ewe lambs show a lower ovulation rate than ewes (Bodin et al., 2007; Beck et al., 1996). The expected ovulation rate is decreased in not yet fully mature ewes by the factor 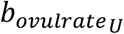 (Fig. 11b). The standard deviation of the ovulation rate depends on a standard deviation target (*ovulrate*_*sd*_^*^) and is changed by *PropRes* times the factor 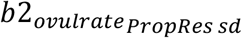 (for fat ewes if 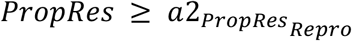). Negative ovulation numbers are set to 0.

**Fig. 11:**
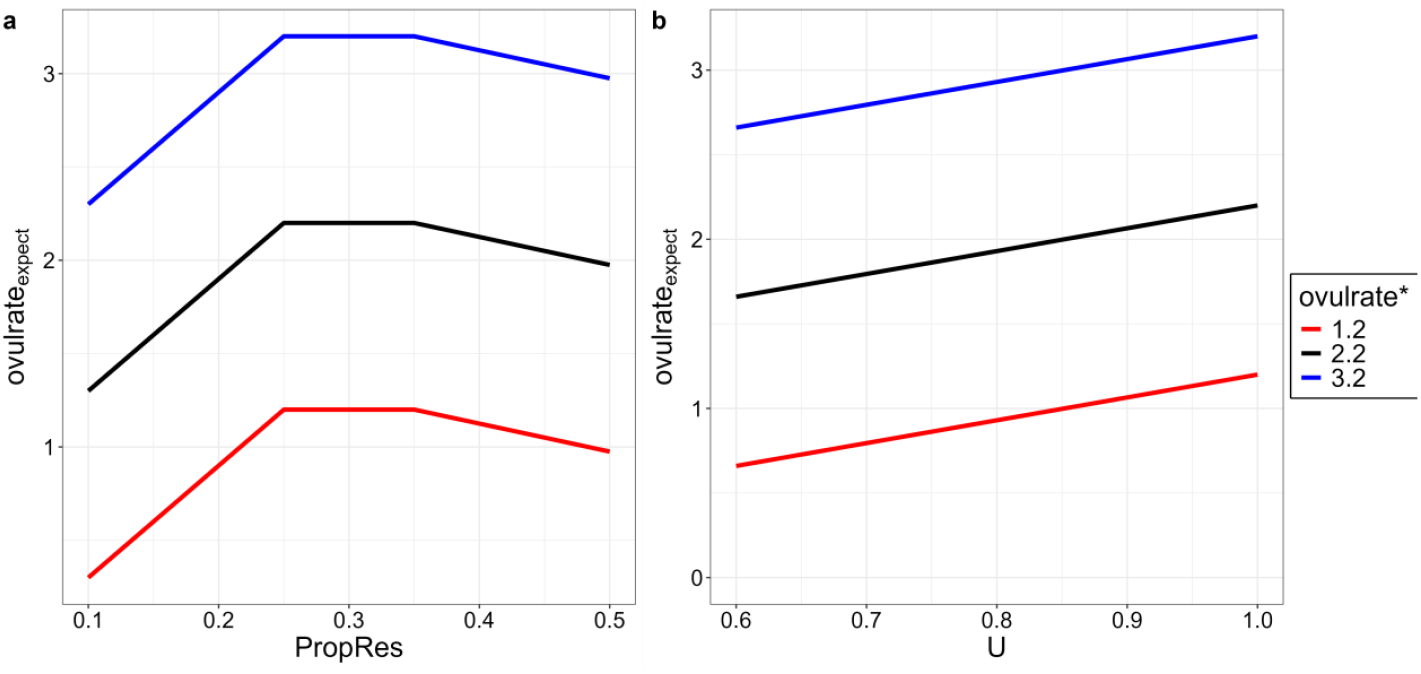
Effect of body reserve level (*PropRes*) and maturity (*U*) on the expected ovulation rate (*ovulrate*_*expect*_). **a** The *ovulrate*_*expect*_ is a function of *PropRes*, the target ovulation rate *ovulrate*^*^, the slope 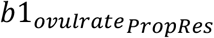 (here −0.6) below the lower *PropRes* limit 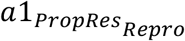 (here 0.25) and the slope 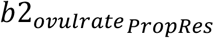 (here −0.15) above the upper *PropRes* limit 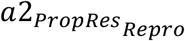 (here 0.35). Maturity *U* was fixed at 1 and the reserve change *deltaPropRes* at 0. **b** The *ovulrate*_*expect*_ as a function of *U, ovulrate*^*^ and the slope 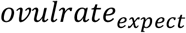 (here −1.35). *PropRes* was fixed at 0.3 and the reserve change *deltaPropRes* at 0.

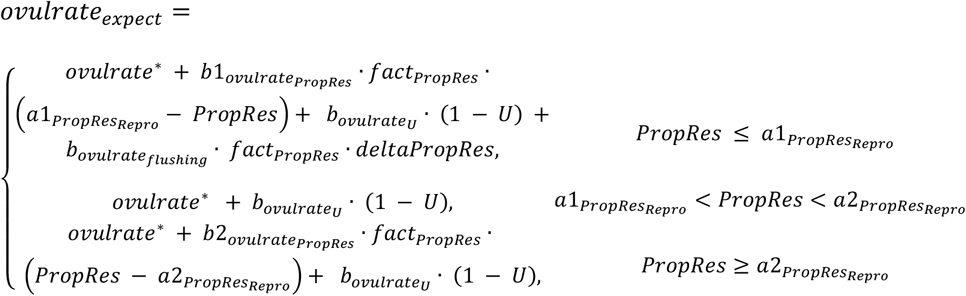

With

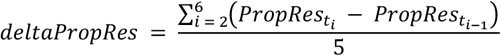

###### Fertilisation, embryonic survival and start of pregnancy

Fertilisation of the ova is possible if the day of the ovulation is within the mating period (*Mating*_*day year*_) and the ewes have reached the minimal age to be mated (*age* ≥ *Agemin*_*mating*_). The number of fertilised ova (*nfertilovum*) is sampled from the ovulation rate (*novul*). For each ovum, fertilisation is independently randomly sampled from a binomial distribution. The expected fertilisation probability depends on the maximum fertilisation probability 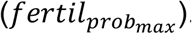.Low and high body fat reserve levels affect the fertilisation rate (Gunn et al., 1972). The expected fertilisation rate is reduced by the factor 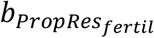 if the *PropRes* is below 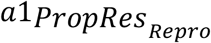 or above 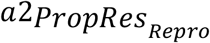.

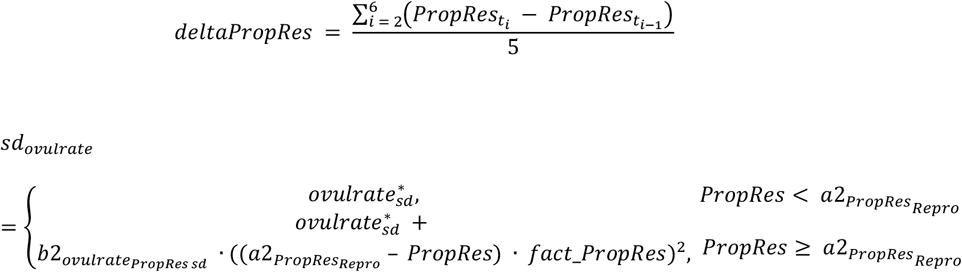

with *X*_*ovul*_ ∼ *B*(*novul, fertil*_*prob*_)

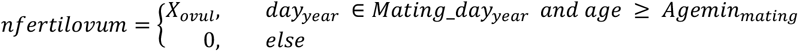

Embryonic survival is randomly sampled from a binomial distribution independently for each fertilised ovum (Michels et al., 1998) according to the expected embryonic survival rate dependent on the number of fertilised ova (1, 2 or ≥ 3) (Diskin and Morris, 2008; Geisler et al., 1977). The number of embryos (*nembryo*) is given by:

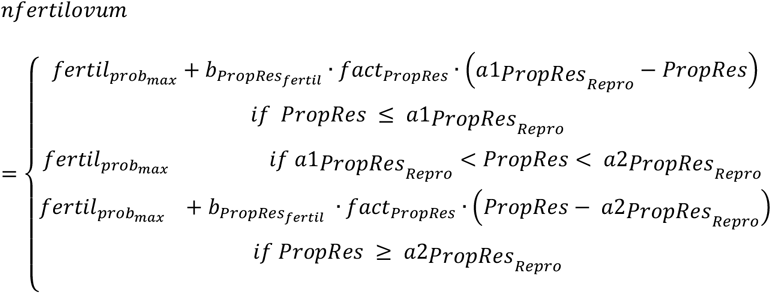

with *X*_*embryo*_ ∼ *B*(*nfertilovum, embryo*_*prob*_)

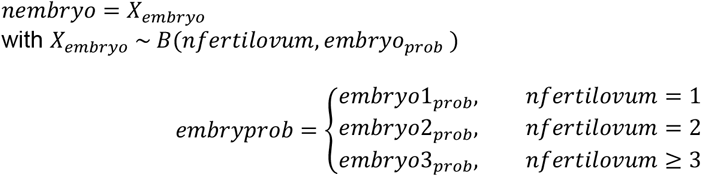

The events include fertilisation and early embryonic death, but occur in the model all on the day of the oestrus and ovulation. If at least one embryo is present, pregnancy starts. No embryonic and fetal losses can occur during the gestation. The number of embryos translates into number of lambs during pregnancy and the number of lambs born (*NLB*).

###### 7.2.2 Birth

At the end of the gestation period, when *Gest*_*time*_ reaches the gestation length *Gest*_*length*_ (Forbes, 1967), gestation ends (*Gest*_*stat*_ is changed from 1 to 0) and the mass of the mother’s gravid uterus (*MassGU*) is translated into lamb birth mass (*MassEmpty*_*Birth*_) with the conversion factor *k*_*MassGULamb*_. The proportion of this mass attributed to each lamb (*PropMassbirth*) is randomly sampled from a normal distribution of mean 1*/NLB* while limiting extreme cases:

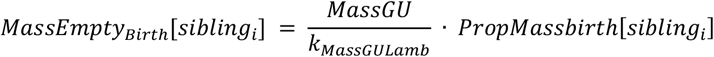

Based on *MassEmpty*_*Birth*_ and lipid reserve proportion at birth (*ResLipProp*_*birth*), initial structural mass (*MassStruct*_*Ini*_), protein reserves (*ResProt*), lipid reserves (*ResLip*) are calculated. Lamb’s sex is randomly sampled from a binomial distribution (0 = female, 1 = male). Birth event triggers the start of lactation (*Lact*_*stat*_ is changed from 0 to 1). The number of lambs suckled (*NLS*) is first inherited from the number of lambs pregnant with (*NLB*) which is set to 0. After the day of birth, *NLS* of the ewe is equal the number of her living lambs that are not weaned yet.

##### 7.3 Survival sub-model

Survival is defined in general and in two particular cases: at birth for lambs and at lambing for the ewe.

###### General survival

At any *age* > 0, general survival is sampled each day from a binomial sampling for each individual who is still alive and with some reserves:

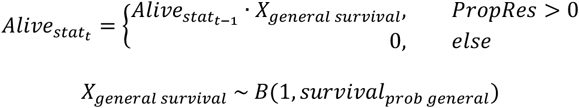

General survival probability (*survival*_*prob general*_) is determined from a Gompertz mortality risk function. It includes a basal mortality rate (*b*_*mort*_), an adjusted effect of body reserves (*deviationPropRes*), and an effect of age (Fig. 12). It is defined as follows:

**Fig. 12:**
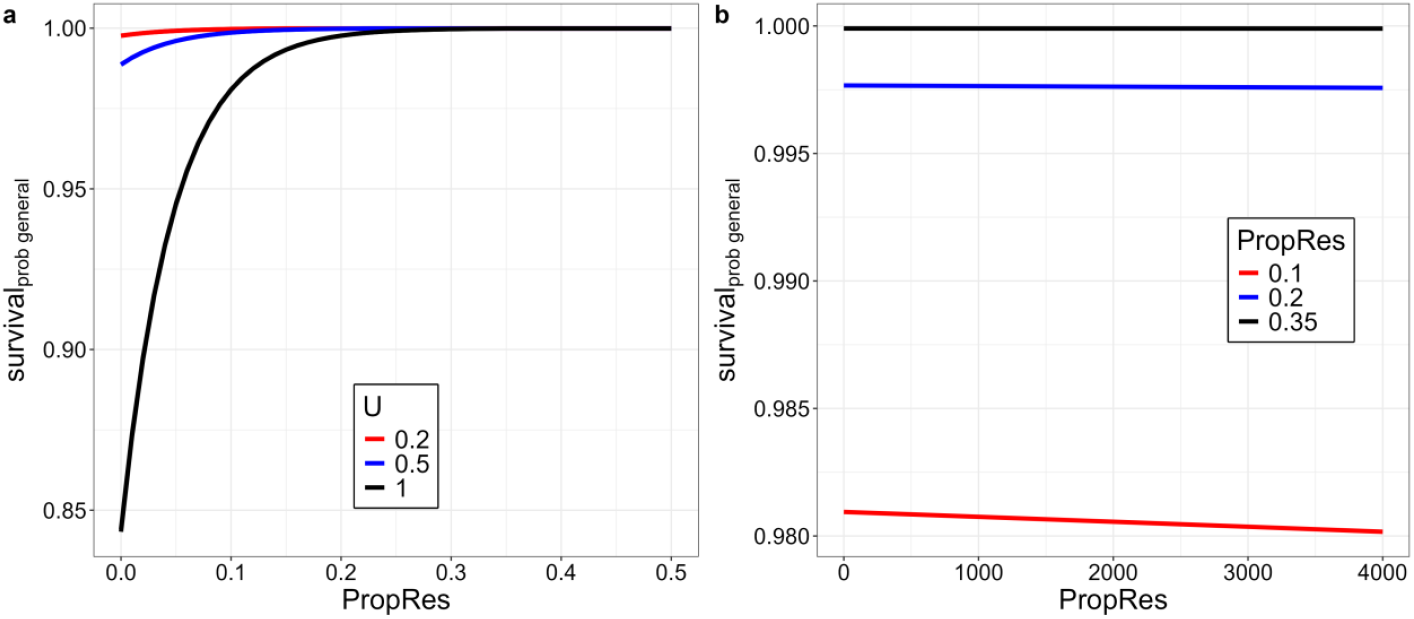
Effect of body reserves (*PropRes*) and *age* on general survival probability (*survival*_*prob general*_). **a** Survival probability as a function of *PropRes* and *U* (with *Age* kept constant at 200 days). **b** Survival probability as a function of age and *PropRes* (with *U* kept constant at 1). Note the different y-axis scales.

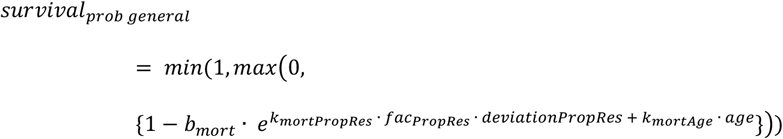

with *k*_*mortPropRes*_ and *k*_*mortAge*_ the coefficients associated with body reserves and age effect, respectively. The variable *deviationPropRes* is defined as the deviation of PropRes from an expected level PropRes adjusted for maturity (*PropRespredU*):

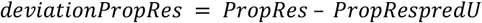

where *PropRespredU* increases from *PropRes*_*birth*_ to *PropRes*_*mature*_ when *U* increases from 0 to 1:

*Neonatal survival:* at birth (*age* = 0) neonatal survival is sampled from a binomial sampling:

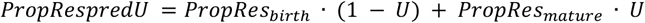

With *X*_*neonatal survival*_ ∼ *B*(1, *survival*_*prob neonatal*_)

Neonatal survival probability is dependent on lamb’s birth mass (*MassEmpty*_*birth*_) with an optimum around medium birth masses (Fig. 13), but decreases at low values due to hypothermia-starvation and at high values due to dystocia (Dwyer et al., 2016; Corbière et al., 2012; Oldham et al., 2011; Scales et al., 1986). Of note, the proposed default parameter values are suitable for average sized breeds and would need to be adjusted for very small or very large sized breeds.

**Fig. 13:**
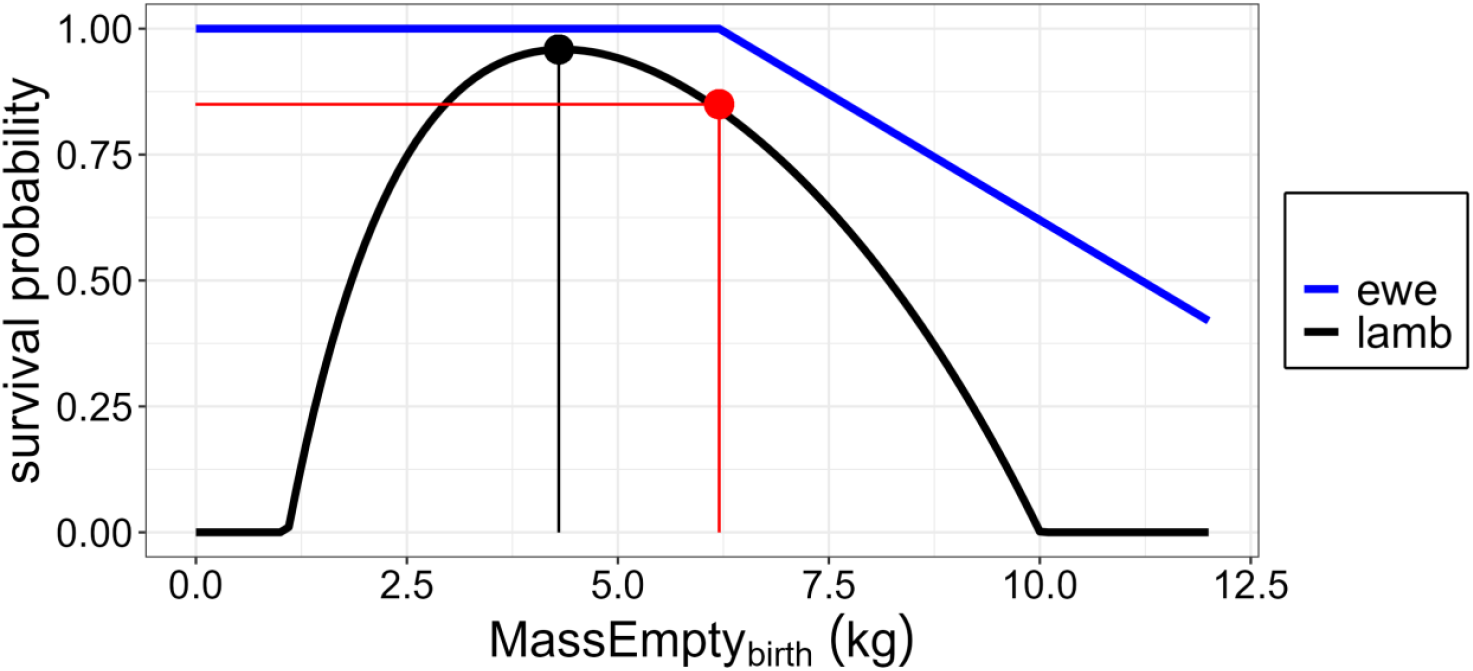
Survival probability of lamb (*survival*_*prob neonatal*_) and its mother (*survival*_*prob moth birth*_) at birth as a function of lamb birth mass (*MassEmpty*_*birth*_). Survival probability of the mother (*survival*_*prob moth birth*_) is determined by the survival probability function of the lamb. *LambMass*_*threshold*_ (red point) is the lamb mass that is higher than the lamb mass with the highest survival probability (black point) and where *survival*_*prob neonatal*_ is equal to *a*_*surv dystocia*_. Above this threshold, *survival*_*prob moth birth*_ linearly decreases with lamb birth mass with the slope *b*_*surv dystocia*_. Parameters used: *a*_*surv dystocia*_ = 0.85 and b_surv dystocia_ = 0.10.

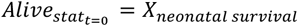

*Ewe survival at parturition:* In addition to general survival, another risk of ewe death of age t is considered at parturition (i.e. when *Lact*_*time*_ = 0):

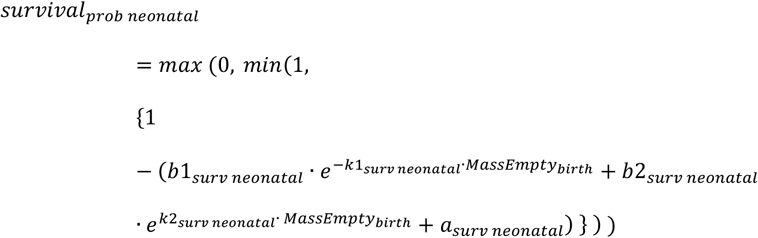

with *X*_*moth survival*_ ∼ *B*(1, *survival*_*prob moth birth*_)

Ewe survival probability at parturition (*survival*_*prob moth birth*_) is assumed to depend on lamb survival and mass at birth. Specifically, if at least one lamb of the ewe dies at birth and has a birth mass (*MassEmpty*_*Birth*_) above a threshold (*LambMass*_*threshold*_), survival probability of the mother is reduced proportionally to the lamb mass due to dystocia with the slope *b*_*surv dystocia*_ (Fig. 13):

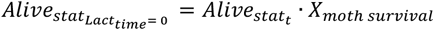

where

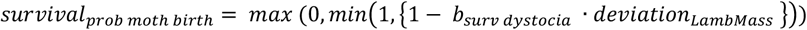

*deviation*_*LambMass*_ = *max*(0, (*MassEmpty*_*Birth*_[*dead lamb*] − *LambMass*_*threshold*_)) Lamb mass threshold above which ewe survival probability decreases (*LambMass*_*threshold*_) corresponds to the mass above the maximum of *survival*_*prob neonatal*_ (cf. black line in Fig. 13) where *survival*_*prob neonatal*_ equals a probability *a*_*surv dystocia*_ (cf. red line in Fig. 13).

###### Slaughter or culling

Individuals can be voluntarily removed (e.g. slaughter or culling) by forcing *Alivestat* to 0 when some criteria are met. For instance, when simulating lifetime trajectories of ewes, lambs produced during the simulation can be systematically removed when reaching the age at weaning to focus on initial individuals.

### Model application

To illustrate how individual variation in model outputs can be simulated, we tested the effects of varying 5 parameters controlling energy allocation trade-offs between the ewe and her lambs. This included 4 input parameters controlling the amount of energy allocated to the offspring during pregnancy and lactation: 2 in the acquisition process (*b*_*AcqPreg*_ and *b*_*AcqLact*_) and 2 in the allocation process (*AllocPreg*^*^ and *AllocLact*^*^). Moreover, a general parameter controlling acquisition was also considered (*b*_*AcqStruct*_) as it should impact the total amount of energy available to the ewe for her own needs (maintenance, growth and reserves) vs. for her lambs (pregnancy and lactation). Individual variation could be considered in many other input parameters however the objective here was to simply illustrate individual variation in ewe-lamb trade-offs, not to implement a full sensitivity analysis. Except the 5 previous input parameters, other parameters were set to the default values as indicated in Table 2. Each one of the 5 input parameters that we tested could take two values (i.e. the low and high values reported in Table 2), resulting in 2^5^ = 32 different parameter combinations. The 32 types and the type with default values (Table 2), were replicated 20 times (by defining 20 initial ewes for each type in the initialisation step). In all cases, no interaction was assumed among individual types or among replicates.

Simulation length (*TSIM*) was set at 4000 days (about 11 years) and results were saved every *dT* = 7 days. The simulation started at day 1 (1^st^ of January) of year 1. The 20 initial ewes were born as orphans in spring (day 85 to 100) of the first year of simulation and suckled artificially *ad libitum* until weaning at 84 days. Feed metabolizable energy content (*MEC*) was derived from a sinus curve with period length 365 days, starting at day 1 of the year. The maximum 11.5 MJ ME / kg DM was reached at day 91 of the year (early spring) and the minimum 7.5 MJ ME / kg DM at day 274 of the year. The feed quantity was not restricted, feed was available *ad libitum* for all animals. Minimal age at mating was set to 8 months (240 days). The mating period took place in winter (between day 300 to 340), resulting in a lambing period in spring. Ewes suckled their lambs for about 3 months (84 days). Suckling and weaned lambs had the same solid feed available as the ewes. The lambs were slaughtered around 4.5 months (at 140 days of age) and thus removed from the simulation. Ewes lived until they died or until the end of simulation.

Simulations were run in R version 4.4.1 (2024-06-14 ucrt) using R Studio (version 2024.09.1) in Windows 11 Pro (version 23H2) on an Intel(R) Core(TM) i7-8665U CPU @ 1.90GHz 2.11 GHz with 32.0 GB RAM, a 64-bit operating system, and a x64-based processor. The model code requires the R packages “geosphere” (Hijmans et al., 2024), “plyr” (Wickham, 2023), “dplyr” (Wickham et al., 2023) and “data.table” (Barrett et al., 2025). The model runtime was recorded with the R package “tictoc” (Izrailev, 2024). The code needed to run the model is available at https://doi.org/10.6084/m9.figshare.28840367.v3.

### Model output analysis

We calculated summary output traits out of the direct model outputs. The summary output traits reflect the mean trait values of the 20 initial ewes (replicates) and their lambs under consideration of factors such as age, parity, ewe age (yearling vs. older), physiological stage and litter size. The ranges of trait values were derived from the default input parameter run and the runs with 5 varying acquisition and allocation parameters.

Repeatability of model outputs was estimated with the R package rptR version 0.9.22 (Stoffel et al., 2017, 2019) using the input parameter combinations as a random grouping factor (1 default input parameter run + 32 varying input parameter runs = 33 groups). The repeatability *R* is the fraction of variance among the groups *V*_*ind*_ out of the total phenotypic variance *V*_*P*_, which is the sum of the variance among the groups *V*_*ind*_ and the within-group residual variance *V*_*res*_ (*R* = *V*_*ind*_*/ V*_*P*_ = *V*_*ind*_ */* (*V*_*ind*_ *+ V*_*res*_)). A highly repeatable trait thus indicates low variation within individual types (e.g. among replicates of a type) relatively to the variation among individual types. Variation among replicates can result from different trajectories of responses due to stochasticity in reproductive and survival events. To take into account this source of variation, an adjusted repeatability (*R*_*adj*_) was estimated by adjusting *R* for two fixed effects: litter size and ewe age (yearling vs. older). The 95 % confidence interval was estimated using 100 parametric bootstraps.

## Results

The model run relatively fast (about 25s). The R scripts to implement the full factorial design and reproduce the simulation outputs are available at https://doi.org/10.6084/m9.figshare.28840367.v3 (sheep_model-main > application > sheepModel_full_factorial_design.R).

### Output traits

The ewe probability to survive until the end of simulation (11 years) was 32% (range min-max = [0, 70]). The 32 ewe types plus the “default” ewe type had on average 5.87 parities [3.63, 8.56]). On average they weaned 9.9 lambs [5.8, 14.2]. Mean litter size at birth over all parities was 1.68 [1.55, 1.86]. Mean lamb birth mass was 5.35 kg [3.9, 7.2]. Neonatal survival was of 84.5% [61, 96]. During the first week of life, lamb daily milk intake was on average 1.26 kg [0.73, 1.87] (assuming a constant milk energy content of 4.7 MJ/kg). At 6 weeks, lamb mass was 15.9 kg [11.9, 21.2]. At weaning, lambs reached a mean mass of 28.1 kg [21.4, 36.2]. Average daily gain from birth to weaning was 275 g/day [196, 372].

Litter size affected a number lamb traits, including birth mass (*NLB* = 1, 2, and 3: 7.57, 4.86 and 3.55 kg), milk intake (e.g., during first week: 1.6, 1.2 and 0.9 kg), lamb mass (e.g. at weaning: 34.1, 27.3 and 22.3 kg) and average daily gain from birth (e.g. at weaning: 315, 267, 223 g/day). Litter size also had a large effect on ewe body mass at lambing (61.8, 58.2, and 55.9 kg) and *PropRes* (0.35, 0.31 and 0.28). Those differences in ewe body mass and reserves progressively reduced during lactation and were almost cancelled at weaning.

### Visualization of trajectories of trait responses

For a given ewe type (defined by the input parameters), diverse lifetime trajectories can occur as a result of different management practices or stochasticity in death and reproduction events. For traits such as *MassTota*l and *PropRes* changes are also particularly affected by seasonal variation in feed energy density. As an example, variation in trajectories is shown in relation to age at first mating (at 1 year (Fig. 14a-b) vs. 2-year of age (Fig. 14c-d)). Lamb growth is also presented at each parity (a singleton in all parities, with a neonatal death in parity 2 in Fig. 14a-b).

**Fig. 14.**
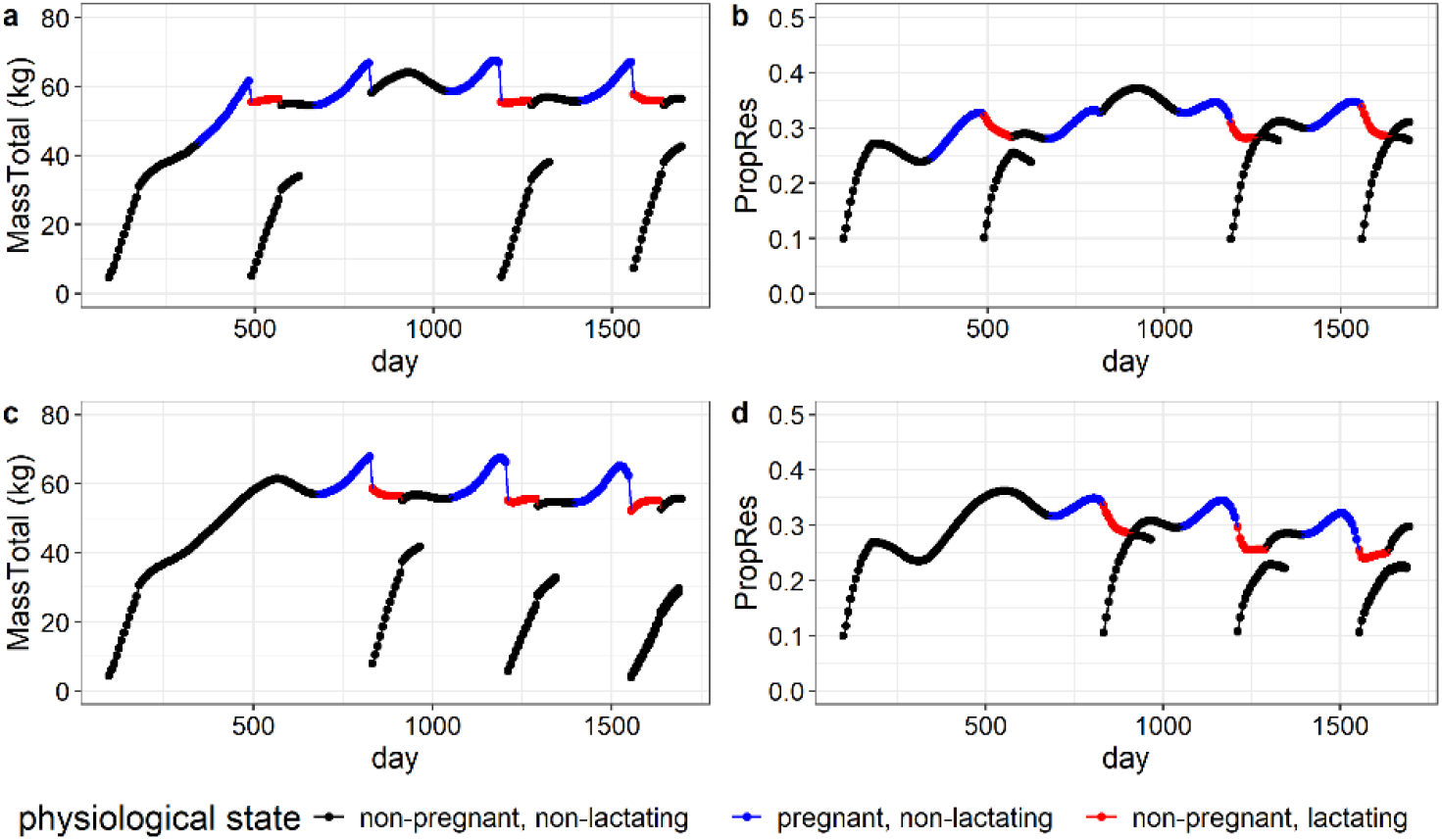
Changes in body mass (*MassTotal*) and body condition (*PropRes*) in a ewe and her lamb depending on age at first parity. The ewe shown in the upper row (**a** and **b**) lambed the first time as yearling, while the ewe shown in the lower row (**c** and **d**) lambed the first time as 2-year-old. Results are presented for 1700 days of simulation with default input parameter.

Lamb growth is affected by litter size at birth and from birth to weaning (Fig. 15). An increase in litter size at birth leads to a decrease in birth mass. During lactation, an increase in suckling litter size leads to an increase in total milk energy provided by the mother (Fig. 15a). In general, this increase of maternal milk production does not compensate the increase in milk energy demand with litter size, so individual lamb milk intake is reduced in large litters (Fig. 15b). Due to low birth mass and competition for milk energy, lambs born and raised in large litters growth slower than others (Fig. 15c). The effect of litter size on lamb growth then decreases when lambs start to eat solid feed (Fig. 15d).

**Fig. 15.**
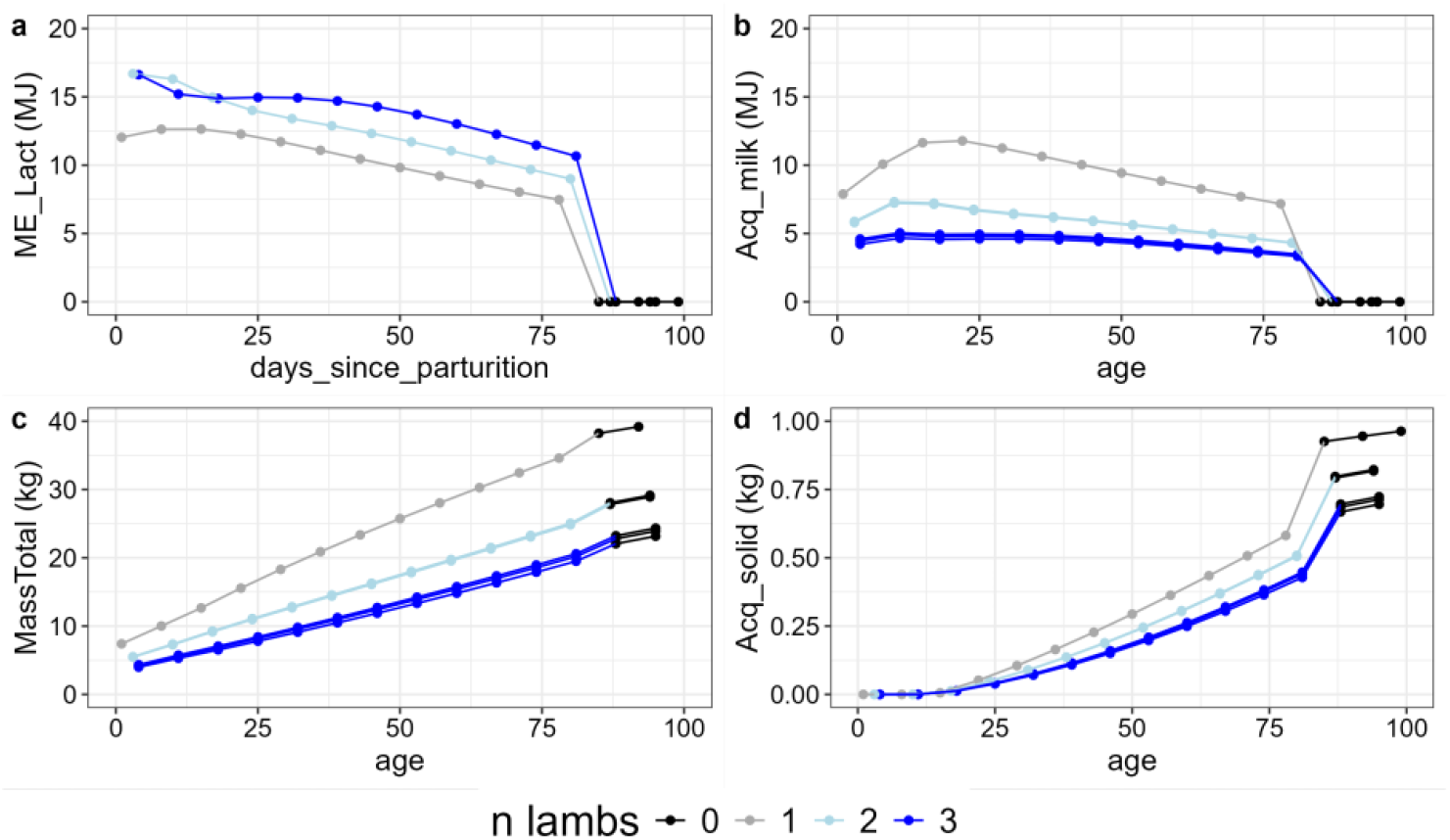
Effect of litter size at birth and during the suckling period on (a) maternal milk energy (*ME*_*Lact*_), (b) individual milk energy intake of each lamb (*Acq*_*milk*_), (c) individual lamb body mass (*MassTotal*), and (d) individual lamb solid feed energy intake (*Acq*_*solid*_). Simulation results of 3 multiparous ewes with default input parameters.

The effect of litter size on maternal traits (body mass and condition, feed intake) around lambing is shown for primiparous and multiparous ewes (Fig. 16).

**Fig. 16.**
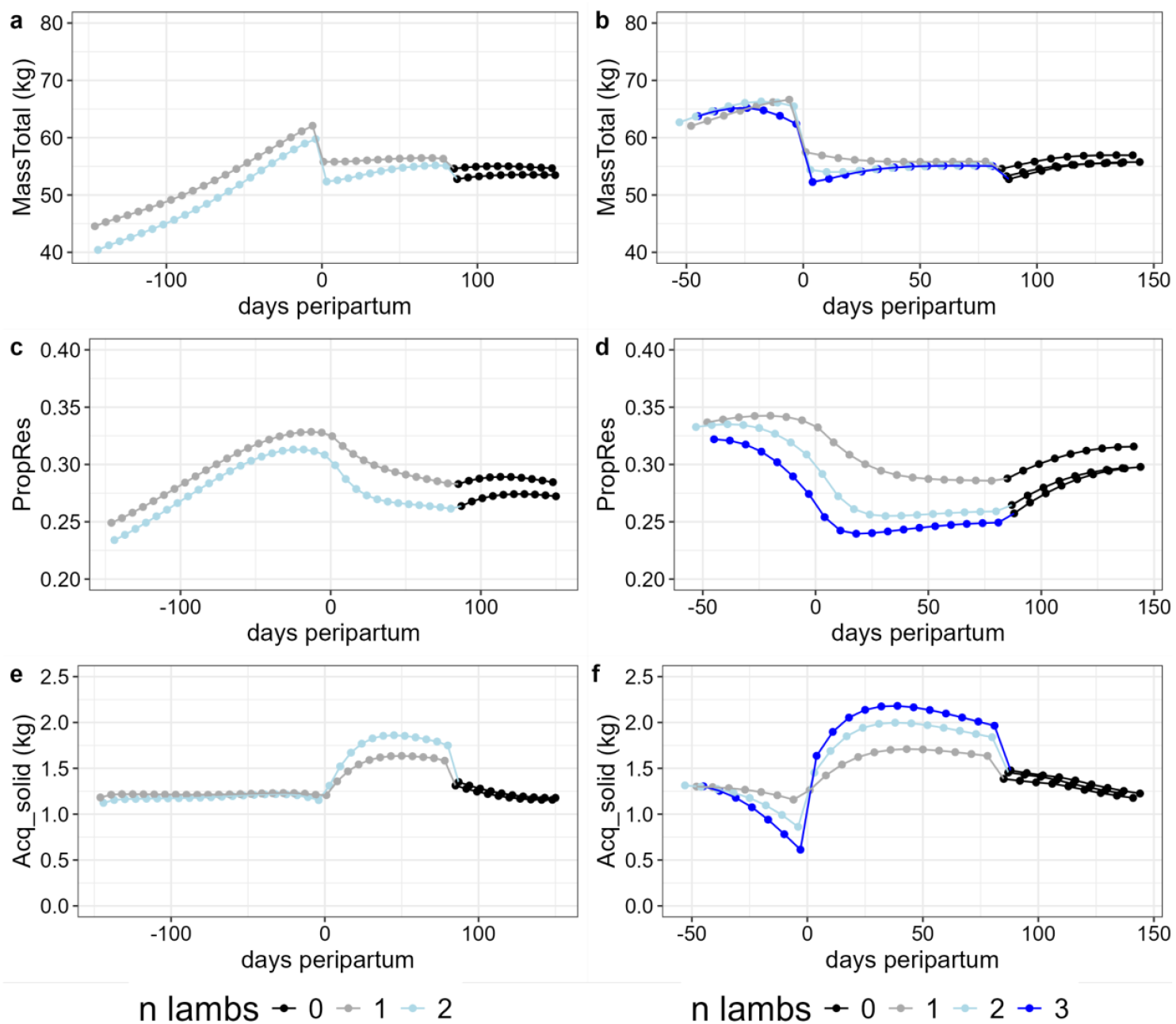
Effect of litter size on maternal traits around lambing in primiparous (a, c, e) and multiparous (b, d, f) ewes. Maternal traits include (a, b) body mass (*MassTotal*), (c, d) body condition (*PropRes*), and (e, f) feed intake (*Acq*_*solid*_). Simulation results of 2 primiparous and 3 multiparous initial ewes with default input parameters.

### Individual variation

We simulated individual variation by testing two different values in five input parameters. This led to 32 parameter combinations that can be considered as different animal “types” (or “genotypes” if input parameters were completely genetically determined). As for the “default” type, each one of the 32 types was replicated 20 times. Those simulations led to additional variation in outputs traits both for the ewe and her lambs. As an illustration, the effects of each input parameter on the means of various maternal and growth traits are shown at weaning (Table 3).

**Table 3:** Effect of the five input parameters on the mean values of model outputs traits observed at weaning. In addition to the individual type run with default values, 32 animal types were run. For each input parameter half a low value (n = 16) and the other had a high value (n = 16). See Table 2 for parameter values.

| Output trait | Default value | Input parameter |  |  |  |  |  |  |  |  |  |
| --- | --- | --- | --- | --- | --- | --- | --- | --- | --- | --- | --- |
| | | $b_{AcqStruct}$ | | $b_{AcqPreg}$ | | $b_{AcqLact}$ | | $AllocPreg^*$ | | $AllocLact^*$ | |
|  |  | low | high | low | high | low | high | low | high | low | high |
| Lamb traits |  |  |  |  |  |  |  |  |  |  |  |
| $MassTotal$ (kg) | 26.1 | 23.6 | 29.0 | 25.9 | 26.7 | 25.0 | 27.4 | 26.3 | 26.1 | 23.5 | 29.0 |
| $PropRes$ (unitless) | 0.24 | 0.22 | 0.25 | 0.24 | 0.23 | 0.22 | 0.25 | 0.25 | 0.22 | 0.21 | 0.26 |
| $Acq_{milk}$ (MJ/day) | 4.82 | 4.08 | 5.04 | 4.57 | 4.50 | 4.09 | 4.98 | 4.72 | 4.35 | 3.60 | 5.47 |
| $Acq_{solid}$ (DM/day) | 0.51 | 0.49 | 0.64 | 0.56 | 0.58 | 0.56 | 0.57 | 0.55 | 0.59 | 0.56 | 0.58 |
| Ewe traits |  |  |  |  |  |  |  |  |  |  |  |
| $MassTotal$ (kg) | 55.2 | 56.2 | 59.4 | 57.8 | 60.0 | 56.5 | 59.3 | 58.4 | 57.5 | 60.1 | 55.7 |
| $PropRes$ (unitless) | 0.26 | 0.28 | 0.30 | 0.29 | 0.29 | 0.29 | 0.30 | 0.25 | 0.22 | 0.33 | 0.26 |
| $Acq_{solid}$ (DM/day) | 1.80 | 1.70 | 1.91 | 1.81 | 1.80 | 1.70 | 1.91 | 1.77 | 1.83 | 1.67 | 1.93 |
| $ME_{Lact}$ (MJ/day) | 8.79 | 7.50 | 9.46 | 8.43 | 8.46 | 7.57 | 9.32 | 8.42 | 8.47 | 6.80 | 10.0 |

*MassTotal* = Body mass; *PropRes* = fat reserve empty body proportion; *Acq<sub>milk</sub>* = energy acquired from milk; *Acq<sub>solid</sub>* = dry matter acquired from solid feed; *ME<sub>Lact</sub>* = milk energy produced by the mother

We then quantified the repeatability *R* of output traits, that is the proportion of output trait variation that can be attributed to variation among the 32 types (the remaining proportion corresponds to the non-repeatable component that can be attributed to variation within each type). Some input parameters had a strong influence on certain output traits but were not changed which led to *R* ≈ 0. For instance, for litter size born *R* = 0.003 (with CI [0, 0.01]) because it was mainly controlled by *ovulrate*^*^ that had a fixed value in our simulations. Moreover, traits that are subject to stochasticity are moderately repeatable. For instance, for the total number of lambs weaned, that is an aggregated trait subject to random variation in reproduction and survival at multiple times during sheep lifetime, *R* = 0.198 [0.10, 0.28]. In contrast, other traits were more strongly determined by input parameters and were more repeatable. For instance, for lamb birth mass *R* = 0.36 [0.24, 0.47]. However higher values were obtained after adjusting R for fixed effects (*R*_*adj*_). For lamb birth mass, *R*_*adj*_ = 0.65 [0.52, 0.72] when adjusted for *NLB, R*_*adj*_ = 0.72 [0.57, 0.80] when adjusted for *NLB* and the age of the mother. For different ewe and lamb traits (*MassTotal, PropRes, Acq*_*solid*_, *ME*_*Acq*_ and *ME*_*Lact*_), values of *R*_*adj*_ were high at different times which shows that those trajectory aspects of our types were highly repeatable (Table 4).

**Table 4:** Adjusted repeatability for lamb and ewe model outputs. Adjusted repeatability estimates *R*_*adj*_ and 95 % confidence interval CI lower and upper boundaries for lamb and ewe model outputs during the suckling period with the input parameter combinations as random grouping effects. The model outputs were adjusted for the number of lambs in the litter and the age of the ewe (yearling vs. older ewe).

| Stage | Birth | Mid-lactation | Weaning |
| --- | --- | --- | --- |
| Lamb traits |  |  |  |
| *MassTotal* | 0.72 [0.57, 0.80] | 0.84 [0.77, 0.90] | 0.88 [0.80, 0.92] |
| *PropRes* |  | 0.94 [0.90, 0.96] | 0.93 [0.88, 0.95] |
| *Acq<sub>milk</sub>* | 0.84 [0.74, 0.90] | 0.82 [0.74, 0.88] | 0.81 [0.73, 0.86] |
| *Acq<sub>solid</sub>* |  | 0.89 [0.81, 0.92] | 0.91 [0.88, 0.95] |
| Ewe traits |  |  |  |
| *MassTotal* | 0.94 [0.90, 0.95] | 0.94 [0.90, 0.95] | 0.94 [0.90, 0.96] |
| *PropRes* | 0.94 [0.90, 0.96] | 0.96 [0.93, 0.97] | 0.97 [0.95, 0.98] |
| *Acq<sub>solid</sub>* | 0.90 [0.83, 0.93] | 0.96 [0.94, 0.97] | 0.97 [0.95, 0.98] |
| *ME<sub>Lact</sub>* | 0.92 [0.87, 0.95] | 0.95 [0.91, 0.97] | 0.95 [0.9, 0.97] |

*MassTotal* = Body mass; *PropRes* = fat reserve empty body proportion; *Acq<sub>milk</sub>* = energy acquired from milk; *Acq<sub>solid</sub>* = dry matter acquired from solid feed; *ME<sub>Lact</sub>* = milk energy produced by the mother

## Author’s point of views

- The main outcome of this article is the presentation of an energy allocation model in sheep that incorporates the energy transfers from the ewe to her lamb(s). The model is presented in detail following the standard ODD protocol of (Grimm et al., 2006, 2020). It represents a tool to (1) simulate individual lifetime trajectories of output traits related to the ewe and her lambs, and (2) implement individual-based simulations studies to investigate the effects of management, feeding or other sources of individual variation (e.g. breeding) on outputs traits, including the potential trade-offs between those traits.
- Compared with previous energy allocation models, our model allows to consider energy allocation trade-offs between the mother and her offspring or among siblings. Our simulations illustrated how those trade-offs can be controlled by various input parameters. Lamb birth mass is determined by the ewe’s allocation to pregnancy while the milk available for the lamb is determined by the ewe’s allocation to lactation. Ewe’s allocation to pregnancy or lactation is acquired from feed intake and potentially from body reserves. Ewe’s allocation to offspring may be at the expense of the amount of energy allocated to her survival and future reproduction (through the storage of body reserves). In contrast, low energy allocation to pregnancy may be at the expense of neonate survival (due to low birth mass) or postnatal growth.
- Model outputs were in a realistic range in general although the prediction of some traits might be refined. For instance, in our simulations adult ewes reached a body mass between 55 and 64 kg, corresponding to a medium size breed. However ewes gave birth to rather heavy lambs (range 3.8 to 7.2 kg on average), especially singleton lambs were quite heavy with a mean birth mass of 7.5 kg. This birth mass is higher than expected from ewes with a similar live weight. For example, adult Romney ewes with a body mass of 61 to 64 kg at mating gave birth to lambs of 5.5 kg for singletons and 4.4 kg for twins (Corner et al., 2013). Scottish Blackface ewes with a mean body weight of 63.7 kg at mating gave birth to lambs of 5.1 to 5.7 kg for singletons and 4.8 to 5.1 kg for twins (Doney et al., 1981). Values of certain parameters might need to be tuned to fit data of a particular breed in particular context. However, the information required for doing this might be challenging to obtain in general. We advise potential users to at least critically assess the range of values observed in the output traits when running a simulation study.
- It is important to bear in mind that some values of input parameters (or certain combinations of those) can lead to values of output traits that are not observed in practice. For instance, when a high value of *AllocPreg*^*^ was combined with high values of intake parameters (e.g. *b*_*AcqStruct*_ = 0.28 and *b*_*AcqPreg*_ = 0.08), singleton lambs were born unrealistically heavy with 10.41 kg on average. In this case, the model predicts an extremely low probability for the ewe and the lamb to survive so it is unlikely that the sheep type with this particular parameter combination actually exists. However other constraints than those assumed in the model may proscribe the existence of particular types. For instance, input parameters may not vary independently from each other because of genetic constraints. When using individual-based model to simulate populations, an absence of correlations among input parameters is commonly assumed but this assumption may not always hold.
- For a certain animal “type”, different trajectories of responses are possible depending on herd management (e.g. age at first mating). The high repeatability of dynamic outputs traits (e.g. body mass, reserves) suggests that those aspects of the trajectories well characterize our different animal types. Those characteristics could be further investigated to analyse the responses of robust ewes, i.e. how they can constantly produce many good lambs despite limited feeding and without compromising their future reproduction and survival. In particular, our model accounts for carry-over effects from one parity to another through the dynamic of body reserves and its effect on fertility and survival.

In conclusion, we provided a detailed description of a new mechanistic model in sheep that incorporates the energy allocation trade-offs between the ewe and her lambs, and among lambs within a litter. Those trade-offs are expected to be central when studying animal robustness in suckling systems. The code of the model associated with this paper provides the building block for implementing individual-based simulations.

## Ethics approval

Not applicable.

## Declaration of generative AI and AI-assisted technologies in the writing process

The authors declare to not have used any generative AI and AI-assisted technologies in the writing process.

## Author contributions

MH: Methodology, Software, Formal analysis, Investigation, Writing – Original Draft, Visualisation, Writing – Review & Editing

RT: Software, Validation, Writing – Review & Editing

FD: Conceptualisation, Methodology, Software, Funding acquisition, Visualisation, Writing – Review & Editing

## Acknowledgements

The authors thank Jérôme Raoul, Nicolas Friggens, Laurence Puillet, Line Saas Kierkegaard and Alban Bouquet for fruitful discussions about mechanistic animal acquisition and allocation models in a genetic context.

